# Drug-induced eRF1 degradation promotes readthrough and reveals a new branch of ribosome quality control

**DOI:** 10.1101/2023.01.31.526456

**Authors:** Lukas-Adrian Gurzeler, Marion Link, Yvonne Ibig, Isabel Schmidt, Olaf Galuba, Julian Schoenbett, Christelle Gasser-Didierlaurant, Christian N. Parker, Xiaohong Mao, Francis Bitsch, Markus Schirle, Philipp Couttet, Frederic Sigoillot, Jana Ziegelmüller, Anne-Christine Uldry, Niko Schmiedeberg, Oliver Mühlemann, Jürgen Reinhardt

## Abstract

Suppression of premature termination codons (PTC) by translational readthrough is a promising strategy to treat a wide variety of severe genetic diseases caused by nonsense mutations. Here, we present two novel and potent readthrough promoters – NVS1.1 and NVS2.1 – that restore substantial levels of functional full-length CFTR and IDUA proteins in disease models for cystic fibrosis and Hurler syndrome, respectively. In contrast to other readthrough promoters that affect stop codon decoding, the NVS compounds stimulate PTC suppression by triggering rapid proteasomal degradation of the translation termination factor eRF1. Our results show that this occurs by trapping eRF1 in the terminating ribosome, causing ribosome stalls and subsequent ribosome collisions, activating a novel branch of the ribosome-associated quality control (RQC) network that involves the translational stress sensor GCN1 and the catalytic activity of the E3 ubiquitin ligases RNF14 and RNF25.

## Introduction

About 10% of all known genetic diseases are caused by nonsense mutations that prematurely truncate the coding sequence (CDS) ^1^. Prototypical and intensively investigated examples are nonsense mutations in CFTR, IDUA, and dystrophin that cause cystic fibrosis (CF), Hurler syndrome (mucopolysaccharidosis type I, MPS I), and Duchenne muscular dystrophy (DMD), respectively ^2-4^. Promoting translational readthrough at such premature termination codons (PTCs) is a promising therapeutic strategy allowing the production of functional full-length protein. Several small molecules that suppress nonsense codon recognition and increase the PTC readthrough frequency have been developed and tested over the past years ^4^. For example, certain types of aminoglycosides, such as geneticin (G-418), induce readthrough of termination codons (TCs) and their potential has been evaluated for different disease models ^5,6^. While clinical trials with aminoglycosides indicated the production of some full-length functional protein and chemically synthesized aminoglycoside variants even showed reduced ototoxicity and kidney liabilities, their clinical application as readthrough drugs is under debate ^3,7^. Multiple screens to identify novel readthrough-promoting compounds resulted in several non-aminoglycoside readthrough promoters, including Ataluren (previously PTC-124 ^8^) and the histamine release inhibiting anti-allergic drug Amlexanox, which was reported to stabilize PTC-containing mRNAs and increase translational readthrough ^9^. Nevertheless, the clinical performance of the currently available readthrough promoters remained modest and controversial, underscoring the need for developing better readthrough promoting drugs.

When an elongating ribosome arrives at one of the three TCs (UAA, UAG, or UGA), a ternary complex consisting of the eukaryotic release factor 1 (eRF1; also known as eukaryotic termination factor 1, ETF1) and the GTP-bound GTPase eRF3 (also known as GSPT1) binds to the aminoacyl-tRNA site (A site). After eRF3-mediated GTP hydrolysis, eRF1 adopts its catalytically active conformation and cleaves the peptidyl-tRNA bond of the peptidyl-tRNA located in the P site of the ribosome ^10-12^. This triggers a rotation between the ribosomal subunits that leads to the ejection of the free polypeptide and of eRF1, ultimately leading to the splitting and recycling of the ribosomal subunits ^13^. In unperturbed cells at normal TCs, translation termination is fast and occurs with high fidelity. Readthrough (i.e. incorporation of a near-cognate aminoacyl tRNA) is observed in <0.1% of termination events and depends on the TC and its surrounding sequence context ^14^. Interestingly and for incompletely understood reasons, readthrough appears to occur more frequently at disease-associated PTCs than at physiological TCs (>1%; ^15^), which opens a pharmacological window for induction of PTC-specific readthrough ^16^. While for most readthrough-promoting agents the mode of action has not yet been determined, aminoglycosides lower the accuracy of ribosomal decoding and thereby increase the probability of a near-cognate tRNA pairing with a TC in the A site, leading to translation beyond the TC ^5^. Alternatively, limiting concentrations of eRF1 and eRF3 also increase the readthrough frequency by kinetically slowing down termination and thus increasing the probability for a near-cognate tRNA binding the A site of ribosomes halted at TCs ^17^. However, depletion of release factors has so far not been exploited as a therapeutical approach to promote readthrough.

Here we report the identification and pharmacological optimization of two small molecule compounds with different chemical scaffolds that promote translational readthrough by causing rapid degradation of eRF1. Both molecules mediate substantial translational readthrough in disease relevant recombinant and primary cell models for CF and Hurler syndrome, respectively. In addition, promising proof of concept studies in a newly engineered MPS I Hurler rat model provide an outlook of their therapeutic potential. Investigations into their mode of action indicates that these compounds trap eRF1 in the A-site of ribosomes, thereby inhibiting translation termination and leading to ribosome collisions. Our work revealed a novel branch of cellular translational quality control that senses and clears the occluded A sites by ubiquitination and rapid proteasomal degradation of eRF1. This pathway requires GCN1 and the two ubiquitin ligases RNF14 and RNF25. The resulting diminished cellular eRF1 concentration explains the increased PTC readthrough observed in cells treated with the compounds

## Results

### High throughput screen identifies novel potent readthrough promoting compounds

To screen for small molecules that promote readthrough at premature termination codons (PTCs), we devised a reporter construct that comprises a 60 bp region of the Cystic Fibrosis Transmembrane Conductance Regulator (CFTR) coding sequence (CDS) with a nonsense mutation at codon 122 (Y122X, amino acid position corresponds to full-length CFTR) fused in-frame between an N-terminal GFP and a C-terminal *Renilla* (RLuc) luciferase (Figure 1A). This readthrough reporter gene was site-specifically integrated into the pre-engineered R4 site of the Jump-In™ HEK293 cell line (hereafter referred to as HEKR4) and constitutively expressed. The resulting HEKR4 PTC reporter cells were used for a high throughput primary screen in which 1.6 × 10^6^ compounds were scored for causing increased RLuc activity. 21’645 compounds increased RLuc activity to >35% of the activity observed with the aminoglycoside Paromomycin, which was used as the positive reference compound (Figure S1A). These hits were further narrowed down using a confirmation screen, which was carried out in triplicates. In addition, a CFTR-WT reporter construct (Figure 1A) stably expressed in HEKR4 cells was used to mark potential false positive hits that increased RLuc activity in a PTC-independent manner. The confirmation and filter screens derived 5198 active hits, 3038 of which were validated by assessing dose-response curves in the RLuc reporter cell model (Figure S1A). The candidates were further tested in HEKR4 cells expressing a version of the human coagulation Factor 9 enzyme harboring a nonsense mutation at amino acid position 29 using an F9 high-content imaging assay format (Figure S1A). At this stage, two scaffolds named NVS1 and NVS2 (Figure 1B) were prioritized and further validated using cellular models for cystic fibrosis (CFTR, Figure 1C) and Hurler syndrome (IDUA, Figure 1D) As shown by confocal imaging of non-permeabilized cells using an anti-S-tag antibody, both NVS1 and NVS2 restored full-length, functional CFTR expression at the cell membrane, with more CFTR being detected from readthrough of the UAA at position 122 and the UAG at position 1282 than of the UGA at position 542 (Figure 1C, upper right). To assess the functionality of the restored full-length protein we conducted a forskolin-stimulated CFTR membrane potential assay after compound treatment. In line with the imaging results, NVS1 and NVS2 promoted readthrough most efficiently on the Y122X mutant, followed by W1282X and G548X, respectively (Figure 1C, lower panels). Notably, both compounds performed better in this assay than the readthrough promoting drug Paromomycin, which served as a positive control. In addition, co-administration of the CFTR inhibitor Inh172 ^18^ abolished the membrane potential in cells expressing CFTR-Y122X, demonstrating that the NVS1 and NVS2 mediated increase of the membrane potential was indeed due to an increase of CFTR activity.

**Figure 1.**
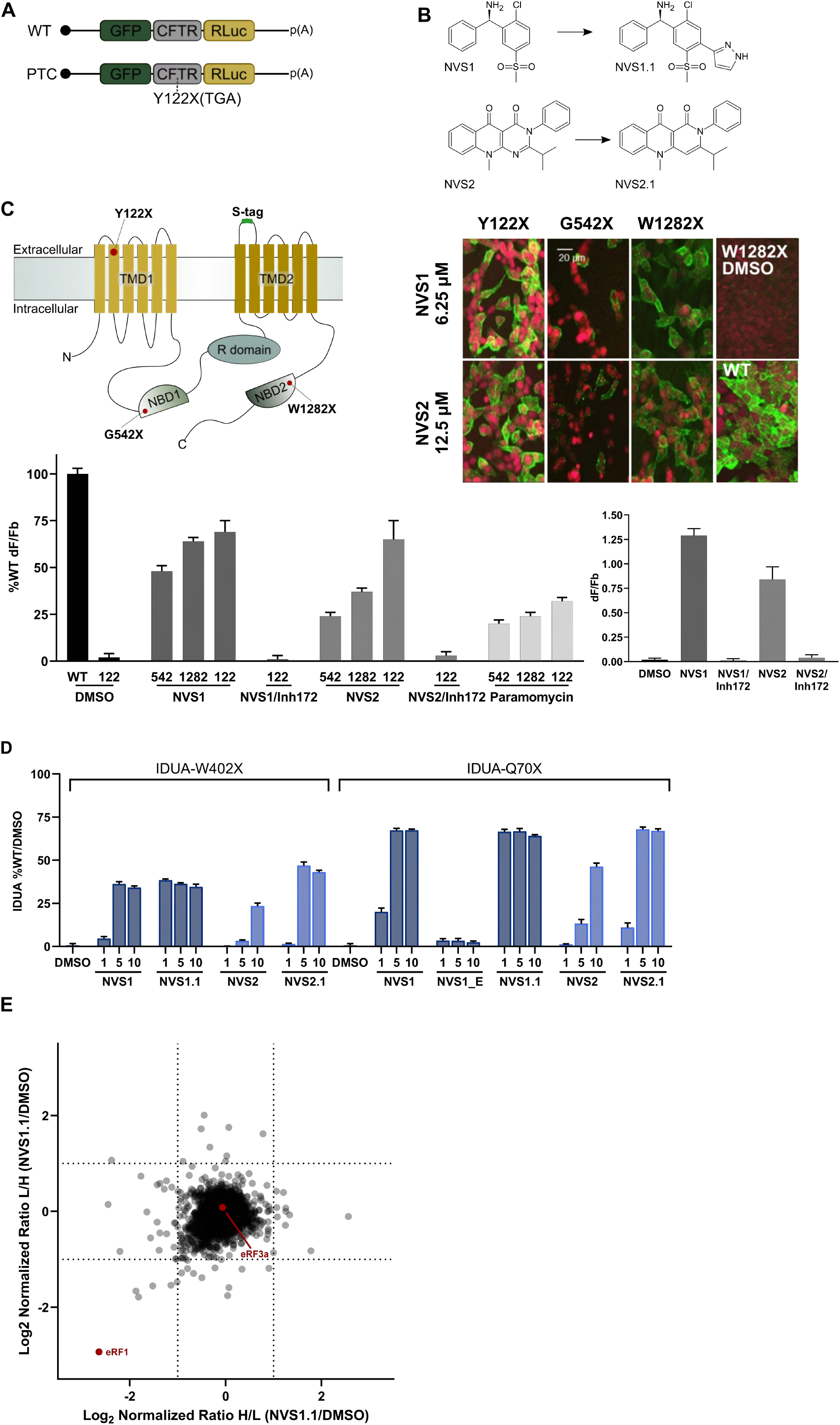
High-throughput screen identifies two small molecules that stimulate PTC readthrough in recombinant cell models. (A) Schematic illustration of the readthrough reporter mRNA used for the high throughput screening. The CDS of GFP, a 60 bp fragment of the CFTR CDS with or without the disease-associated PTC mutation (Y122X), and the RLuc CDS are fused in-frame and depicted by green, grey and yellow boxes, respectively. The 5’ 7-methyl-G cap (black dot) and polyadenylated tail at the 3’end (p(A)) are also shown. (B) Chemical structures of the molecules NVS1 and NVS2 and their pharmacologically optimized derivates NVS1.1 and NVS2.1, respectively. (C) NVS1 and NVS2 restore CFTR expression with nonsense mutations in HEKR4 cells. Upper left: Domain organization of CFTR, depicting transmembrane (TMD), nucleotide-binding (NBD), and regulatory (R) domains and the positions of the tested nonsense mutations Y122X[UAA], G542X[UGA], and W1282X[UAG]. The surface epitope (S-tag) allows detection of correctly inserted CFTR into the cell membrane. Upper right: Cells expressing the indicated CFTR mutants and WT CFTR were treated with 6.25 μM NVS1, 12.5 μM NVS2, or 10 mM Paromomycin for 48 hours. Immunostaining with an anti-S-tag antibody (green) detects active CFTR, nuclei were stained with DRAQ5 (red). Lower left: The CFTR chloride channel activity was assessed by membrane potential measurements in cells expressing the indicated CFTR nonsense mutant pre-treated with 10 μM NVS1, NVS2, or 10 mM Paromomycin, for 48 hours before stimulation with 20 μM Forskolin. Activities are indicated as a percentage of the activity in untreated WT CFTR-expressing cells. Lower right: Membrane potential measurement after addition of 10 μM of the selective CFTR chloride channel blocker 172 (Inh172) for 20 minutes in the presence of NVS1 or NVS2. (D) Restoration of α-L-iduronidase activity in HEKR4 cells expressing IDUA Q70X and W402X mutants treated with 1, 5, or 10 μM of the indicated compounds for 48 hours. NVS1 and NVS2 are compared with their chemistry optimized derivatives NVS1.1 and NVS2.1. NVS1_E is the inactive enantiomer of NVS1. Enzymatic activities were measured and are displayed as percentage of IDUA activity measured in untreated cells expressing WT IDUA. (E) Proteome changes induced by NVS1.1 were monitored by SILAC-MS analysis of HEKR4 cells treated with 1 μM NVS1.1 or DMSO for 24 hours. The experiment was performed twice, once by labeling the NVS1.1 treated cells with the heavy isotope (H) and the DMSO treated cells with the light isotope (L), and once with inverted isotope labeling. The log_2_ of the normalized H/L or L/H ratios for each detected protein is plotted along the x- and y-axis, respectively.

With the aim to apply the readthrough compounds ultimately in a Hurler syndrome rat model, HEKR4 cells expressing IDUA-Q70X and IDUA-W402X were used for structure activity relationship (SAR)guided compound optimization. In total, 179 derivatives of NVS1 and NVS2 were synthesized and compared for their ability to restore IDUA enzyme activity. Of those, the Diphenylmethanamine derivate NVS1.1 and the Pyrimido (4,5-B) Quinoline-4,5 (3H,10H)-Dione NVS2.1 (Figure 1B; patents WO2014/091446A1 and WO2015/186063A1) were selected for further testing (Figure 1D). The chemically optimized molecules NVS1.1 and NVS2.1 outperformed the HTS-derived compounds NVS1 and NVS2 in potency, with superior IDUA activity restoration in both the Q70X and the W402X cell models. Neither DMSO nor treatment with an inactive enantiomer of NVS1 (NVS1_E) restored detectable IDUA enzyme activity. Liquid chromatography followed by mass spectrometry (LC-MS)-based peptide analysis of NVS1.1, NVS2.1, and Paromomycin-treated IDUA-W402X cell lines showed that the amino acid glutamine (Q) was preferentially incorporated at the UAG nonsense codon and tryptophane (W) at the UGA nonsense codon, while UAA led to the incorporation of tyrosine (Y), glutamine (Q) or lysine (K) (Figure S1B).

To further characterize the newly identified and optimized compound NVS1.1, we assessed cellular proteome alterations upon stimulation of translational readthrough. As an unbiased approach, we performed quantitative proteomic profiling using stable isotope labeling by amino acids in cell culture (SILAC) ^19^ in parental HEKR4 cells that were either incubated with NVS1.1 or DMSO as a negative control). The SILAC experiment was repeated with reversed isotope labeling of the two conditions, and the abundance ratio between the NVS1.1-treated and the control sample was determined for each protein detected in both experiments (Figure 1E). Remarkably, we observed a specific 6.5 to 7-fold reduction of eRF1 upon NVS1.1 treatment in both experiments, suggesting that NVS1.1 promotes the rapid and specific degradation of eRF1. Since it has been previously shown that the reduction of translation termination efficiency by the depletion of eukaryotic release factors leads to stop codon suppression ^20-22^, the finding that NVS1.1 causes the proteasomal degradation of eRF1 explains its readthrough-promoting activity. Whereas it was shown that eRF3 degradation results in co-depletion of eRF1 ^20,23^, we observed no effect of NVS1.1 on eRF3 abundance, indicating that eRF3 stability is not regulated by eRF1 levels (Figure 1E).

### NVS1.1 and NVS2.1 restore synthesis of full-length functional protein in human Hurler syndrome primary cell models

To determine the therapeutic potential of the readthrough compounds NVS1.1 and NVS2.1 we extensively tested them in primary fibroblasts derived from Hurler syndrome patients with homozygous mutations in the IDUA gene. Reduced α-L-iduronidase (IDUA) activity, a glycosidase involved in the breakdown of glycosaminoglycans (GAG), leads to a spectrum of disorders collectively referred to as Mucopolysaccharidosis type I (MPS I) and the disease burden correlates with residual α-L-iduronidase amounts ^24,25^. Hurler syndrome represents the most severe form of MPS, which is characterized by complete α-L-iduronidase deficiency and leads to death in early childhood. The clinically most relevant and most frequently found mutations in Hurler syndrome are the homozygous nonsense mutations W402X[UAG] and Q70X[UAG], which were used earlier in recombinant cell models to optimize our readthrough promoters (Figure 1E) ^25^. The patient-derived primary fibroblast cells lack any detectable functional α-L-iduronidase and consequently accumulate large amounts of GAG in an increased number of enlarged lysosomal compartments ^26^. Because it is the rate-limiting enzyme in the lysosomal GAG processing, already small amounts of restored α-L-iduronidase activity can clear the cellular overload of GAG and attenuate disease severity ^27-29^. In our patient cells, treatment for 7 days with NVS1.1 (Figure 2A) or NVS2.1 (Figure S2A) led to a dose-dependent increase of IDUA activity, accompanied by a kinetically similar GAG reduction. Both compounds cleared the cellular GAG load below the assay’s detection limit after 7 days of incubation (Figures 2A and S2A). Based on previous reports showing that loss of function mutations of IDUA lead to upregulation of other genes involved in GAG degradation ^30^, we monitored the activity of the hydrolase -glucuronidase (GUSB). Upon depletion of the cellular GAG levels, we observed a more than twofold reduction of GUSB activity, indicating that the NVS1.1-mediated IDUA restoration has a stabilizing effect on the GAG degradation pathway.

**Figure 2.**
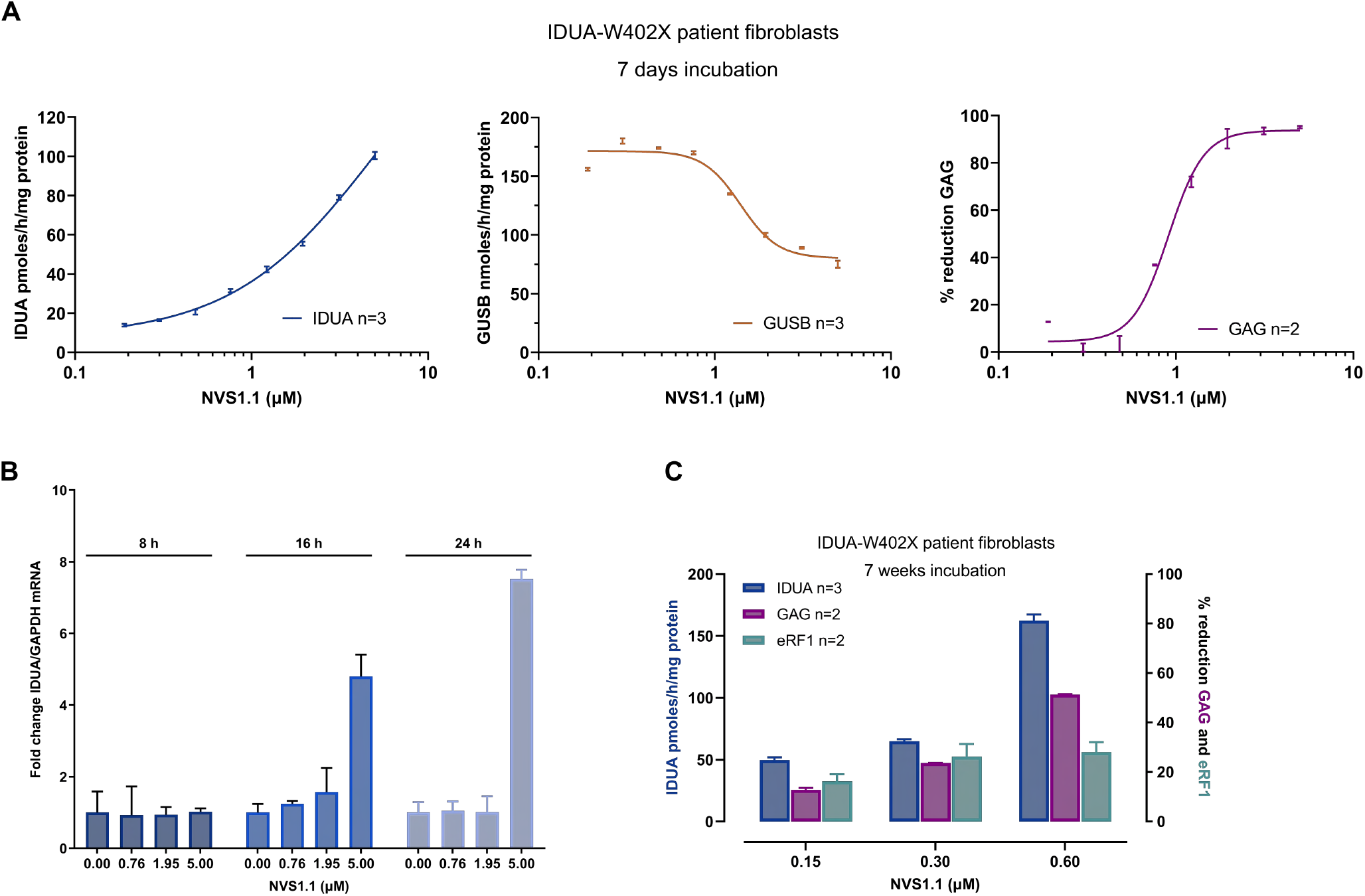
NVS1.1 leads to functional restoration of α-L-iduronidase in Hurler patient fibroblasts. (A) Primary fibroblasts deriving from patients homozygous for the IDUA W402X mutation were treated for 7 days with the indicated concentrations of NVS1.1. Subsequently, α-L-iduronidase (IDUA) and -glucuronidase (GUSB) activities were assessed by measuring the conversions rates of the surrogate substrates 4-Methylumbelliferyl-α-L-iduronide (pmol/h/mg protein) and 4-Methylumbelliferyl-β-D-glucuronide (nmol/h/mg protein), respectively. Total glycosaminoglycan (GAG) levels were determined by a colorimetric assay and normalized to the vehicle-treated control sample. (B) RT-qPCR assay to assess the IDUA-W402X mRNA levels (normalized to GAPDH mRNA) upon treatment of the Hurler primary fibroblasts with different concentrations of NVS1.1 for 8 – 24 hours. (C) Primary IDUA-W402X fibroblasts of Hurler patients were cultured in different concentrations of NVS1.1 with medium and compound exchange every 3^rd^ day. After 7 weeks, the IDUA activity (left y-axis) and total GAG levels (right y-axis) were determined as in (A). The eRF1 protein abundance (right y-axis) was assessed by immunoblot (using beta-actin as a loading control) and is expressed relative to the DMSO control condition.

Since translational readthrough of PTCs can inhibit the degradation of nonsense mRNAs by the nonsense-mediated mRNA decay (NMD) pathway ^31^, we tested if NVS1.1 inhibits NMD of the IDUA-W402X mRNA. RT-qPCR time course experiments in the IDUA-W402X fibroblasts showed that up to 2 μM NVS1.1, IDUA mRNA remained essentially unchanged, while with 5 μM NVS1.1, it increased over time (Figures 2B and S2B). Since we measured increased IDUA activity at NVS1.1 concentrations <1 μM, the observed increase in IDUA protein and enzymatic activity can be attributed primarily to translational readthrough and is not the result of higher mRNA levels due to NMD inhibition.

Since NVS1.1 and NVS2.1 are well tolerated by the patient fibroblasts up to 10 μM, we proceeded to long-term administration of the drugs. In IDUA-Q70X patient fibroblasts, α-L-iduronidase was increased in a dose-dependent manner after 7 weeks of treatment with either of the two compounds (Figure S2C) and comparable levels of IDUA activity were restored by both compounds in IDUA-W402X fibroblasts (Figures 2C and S2D). Importantly, for both compounds the dose-dependent restoration of IDUA activity correlated with the extent of eRF1 depletion and GAG reduction. Treatment with 0.6 μM NVS1.1 reduced GAG, which is the most disease-relevant parameter, to half compared to untreated fibroblasts (Figure 2C) and 2 μM NVS2.1 reduced the accumulated GAG by 80% (Figure S2D). This result indicates that a long-term application of a low dose of NVS1.1 or NVS2.1 might have therapeutic potential for treating Hurler syndrome.

### NVS1.1 is efficacious in a rat Hurler IDUA-W401X animal model

For proof-of-concept studies, we mandated a contract research organization with the engineering of a rat model for Hurler syndrome homozygous for the W401X mutation in the IDUA gene, which mirrors the human disease-linked W402X mutation. To verify responsiveness of the animal model to our compounds, we isolated fibroblasts and treated these primary cells for 7 days with NVS1.1 and NVS2.1. Both compounds restored IDUA enzyme activity and reduced the GAG levels in a dose-dependent manner (Figure 3A and Figure S3A).

**Figure 3.**
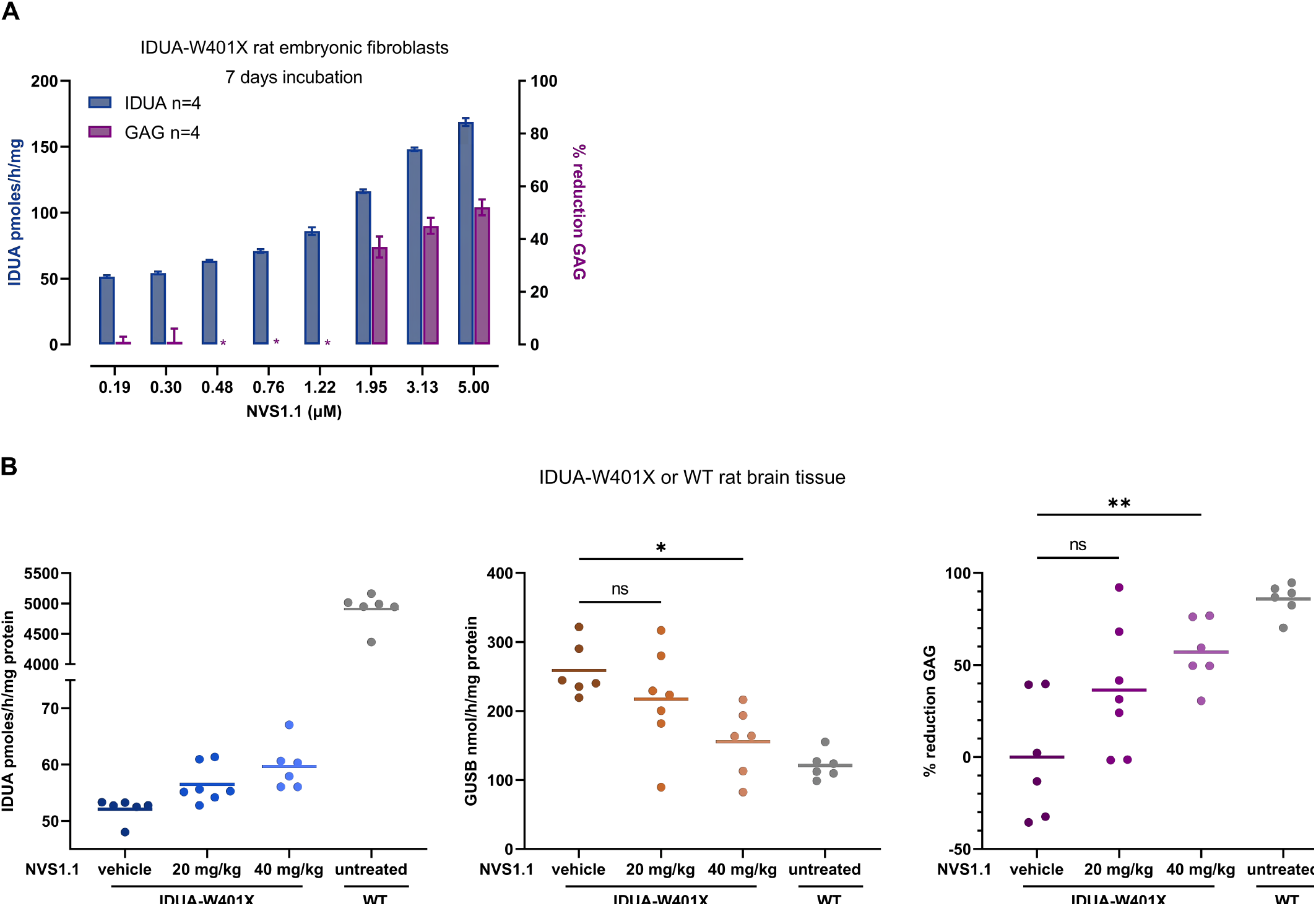
In a rat Hurler disease model, NVS1.1 restores sufficient IDUA enzyme activity to reduce the glucosamine load in the brain. (A) Freshly isolated rat fibroblasts of a Hurler animal model homozygous for IDUA-W401X were treated for 7 days with indicated concentrations of NVS1.1. The IDUA enzyme activity and total GAG levels were determined as in Figure 2A. Asterisks denote GAG measurements below the assay detection limit (resulting in negative values) that were removed from the dataset. (B) Rats homozygous for the IDUA W401X mutation were treated with 20 mg/kg (n=7) or 40 mg/kg (n=6) NVS1.1, or with the vehicle formulation only (n=7) for 14 days. A similar number of untreated animals with WT IDUA was analyzed for comparison. After scarifying the animals, the brain tissue was extracted and the activities of IDUA and GUSB along with the total GAG levels were determined as in Figure 2A. In each graph (IDUA, GUSB, GAG), one dot represents the average of four technical replicates carried out with brain tissue from one animal. *P* values were calculated by one-way ANOVA followed by Dunnet’s multiple comparisons test. *P* values of ≤ 0.05, p≤ 0.01, and >0.05 are depicted as *, **, and ns (non-significant), respectively.

Based on the results obtained with the rat fibroblasts, the IDUA-W401X rats were then used to investigate the *in vivo* efficacy of NVS1.1 and NV2.2. The highly soluble NVS1.1 compound showed good drug-like properties with favorable pharmacokinetics, metabolic stability, and suitability for oral administration. Acceptable profiles for oral *in vivo* pharmacology studies were derived for NVS2.1, too. Furthermore, bioavailability tests showed distribution of NVS1.1 and NVS2.1 into the cerebrospinal fluid (CSF). Brain exposure is a prerequisite for a successful future Hurler syndrome drug, since the current standard of care – mainly enzyme replacement therapy, which does not reach the central nervous system – hardly addresses the early onset and progressing brain defects. Therefore, our initial *in vivo* efficacy studies mainly focused on IDUA restoration and biomarker responses in rat brain tissue. Hurler rats were treated with NVS1.1 or NVS.2.1, and brain tissues were then analyzed for restored IDUA enzyme activity, GUSB enzyme response and total GAG reduction as described earlier for the patient and rat-derived fibroblasts. Daily oral dosing for 14 days with 40 mg/kg body weight NVS1.1 and NV2.1 significantly restored IDUA enzyme activity and reduced the GAG load, whereas the 20 mg/kg dosing for both compounds showed clear but not statistically significant restoration effects (Figures 3B and S3B). Both 40 mg/kg doses of NVS1.1 and NVS2.1 restored approximately 1% of the IDUA activity measured in brain tissue of WT rats (untreated, WT), which resulted in a reduction of GAG by 36% and 57% in the brains of Hurler rats treated with 20 mg/kg and 40 mg/kg NVS1.1 and about 50% with 40 mg/kg NVS2.2, respectively (Figure 3B). Interestingly, as shown earlier for the primary patient fibroblast cell models (Figures 2A and S2A), the GUSB enzyme activity increased more than 2-fold in the Hurler rats compared to WT rats but was dose-dependently reduced by NVS1.1 and NV2.1, reaching approximately 1.3-fold of WT rats with both compounds when dosed at 40 mg/kg (Figure 3B, S3B). Collectively, our data suggests that the pharmacologically optimized molecules NVS1.1 and NVS2.1 restore sufficient amounts of IDUA to clear the cellular GAG levels *in vivo* within two weeks of oral administration in the Hurler syndrome rat model.

### NVS1.1 induces ubiquitination of eRF1 on lysine 279, leading to proteasomal eRF1 degradation

As mentioned above by assessing the changes in the proteome upon treatment with NVS1.1, we identified a specific and substantial depletion of the translation termination factor eRF1 (Figure 1F), providing a mechanistic explanation for the readthrough-promoting activity of the NVS1.1. Canonical protein decay occurs via the proteasome, which recognizes, unfolds, and degrades proteins marked with polyubiquitin. Co-treatment of cells with NVS1.1 and the proteasome inhibitor Bortezomib (BTZ) fully stabilized eRF1, indicating that NVS1.1-induced eRF1 depletion occurs by proteasomal degradation (Figure 4A). To investigate whether NVS1.1 induces the expected ubiquitination of eRF1, we immunoprecipitated eRF1 from cells that were treated with a high dose (25 μM) of NVS1.1 for a brief time (30 min) to capture transient modifications on eRF1 before its degradation. Subsequently, we performed immunoblot analysis with antibodies against the bait protein and ubiquitin (Figure 4B). In the NVS1.1-treated samples, mono-, bi- and tri-ubiquitinated eRF1 was detected, while no eRF1 ubiquitination was observed in the control sample, demonstrating that NVS1.1 triggers ubiquitination of eRF1. The eRF1 immunoprecipitates were also analyzed by label-free mass spectrometry, which showed a strong enrichment of ubiquitin upon treatment of the cells with NVS1.1 (Figure S4). Furthermore, the analysis of peptides with di-glycine remnants conjugated to the epsilon amino group of lysines (K), which allows the identification of ubiquitination sites in a protein ^32^, identified K279 of eRF1 as the main site for NVS1.1-induced ubiquitination (Figure 4C).

**Figure 4.**
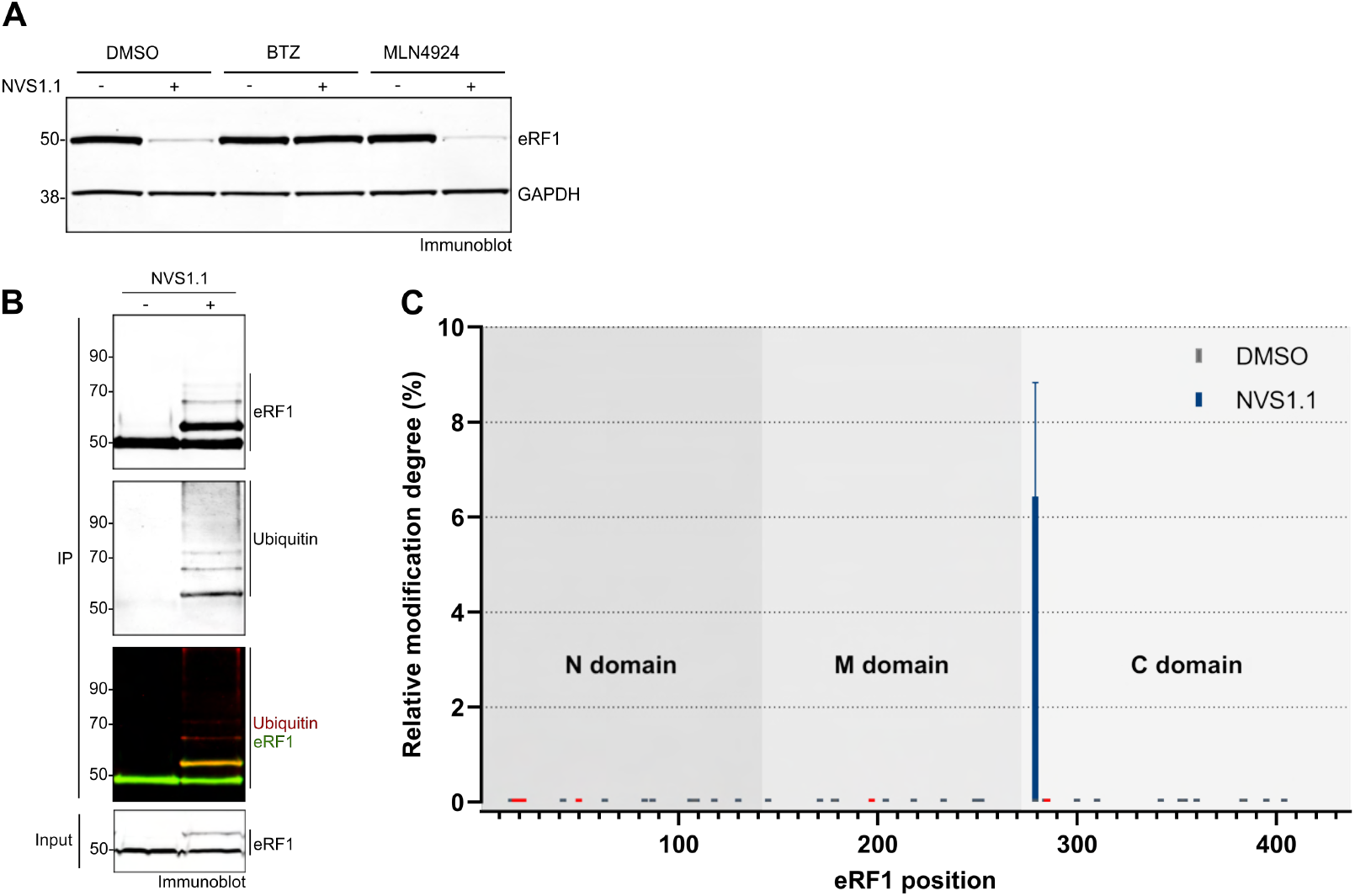
NVS1.1 induces proteasomal degradation of the translation termination factor eRF1 via ubiquitination of K279. (A) Representative immunoblot showing the protein levels of eRF1 in HEKR4 PTC reporter cells when treated with 2.5 μM NVS1.1 for 6 hours in the presence of 0.5 μM of the proteasome inhibitor Bortezomib (BTZ) or 1 μM of the neddylation inhibitor MLN4924 (DMSO was used as control treatment). GAPDH served as a loading control. (B) HeLa cells were treated with 25 μM NVS1.1 for 30 min. Subsequently, eRF1 was enriched by immunoprecipitation, and the eluates were analyzed by immunoblot probing for eRF1 and ubiquitin. (C) Immunoprecipitates from (A) were analyzed by label-free mass spectrometry and the peptides were inspected for glycine-glycine (diGly) remnants on lysine residues originating from ubiquitination events. The bar plot depicts the averaged diGly frequency (y-axis) per lysine residue on the eRF1 protein sequence (isoform1, x-axis) along with its domain organization represented by the background shadings. Grey and blue bars represent lysine residue that were covered by peptides in at least two replicates per condition in the DMSO and NVS1.1 treated samples; lysines depicted in orange were not detected above the threshold.

Since numerous previously identified small molecule degraders act by modulating the receptor specificity of Cullin-RING ligase 4 (CRL4), we addressed if NVS1.1 employs a similar mechanism. CRL4 ligases require neddylation for their activation ^33-35^, which can be prevented by treating the cells with the neddylation inhibitor MLN4924. However, NVS1.1-induced eRF1 depletion was not inhibited by the co-treatment of the cells with MLN4924 (Figure 4A), indicating that NVS1.1 is not a molecular glue degrader that employs CRL4 E3 ligases, but instead has a different mode of action.

### NVS1.1-mediated eRF1 degradation requires the catalytically active E3 ligases RNF14 and RNF25

To identify ubiquitin ligases involved in the NVS1.1-dependent ubiquitination of eRF1 we employed two orthogonal approaches, a genome-wide CRISPR knockout screen scoring for cells that survive under high NVS1.1 concentrations and a genome-wide siRNA screen scoring for reduced readthrough of the *Renilla* luciferase (Rluc) readthrough reporter in cells exposed to NVS1.1. Both screens independently revealed the two E3 ubiquitin-protein ligases RNF14 (Ring Finger Protein 14, UniProt ID Q9UBS8) and RNF25 (Ring Finger Protein 25, UniProt ID Q96BH1) as top hits (Figure 5A). The two proteins contain a RING finger domain typical for E3 ubiquitin ligases, although RNF14 lacks one of the conserved histidines in the RING-type domain 2. Little is known about the function and ubiquitination targets of RNF14 and RNF25. RNF14 has been reported to be a membrane bound E3 ubiquitin ligase that is associated with the androgen receptor and acts as a co-activator of its transcriptional activity ^36,37^. Furthermore, RNF14 is a regulator of TCF/β-catenin-mediated transcription ^38^. RNF25 has been shown to support NF-kappaB-mediated transcription by interacting with its p65 subunit ^39^.

**Figure 5.**
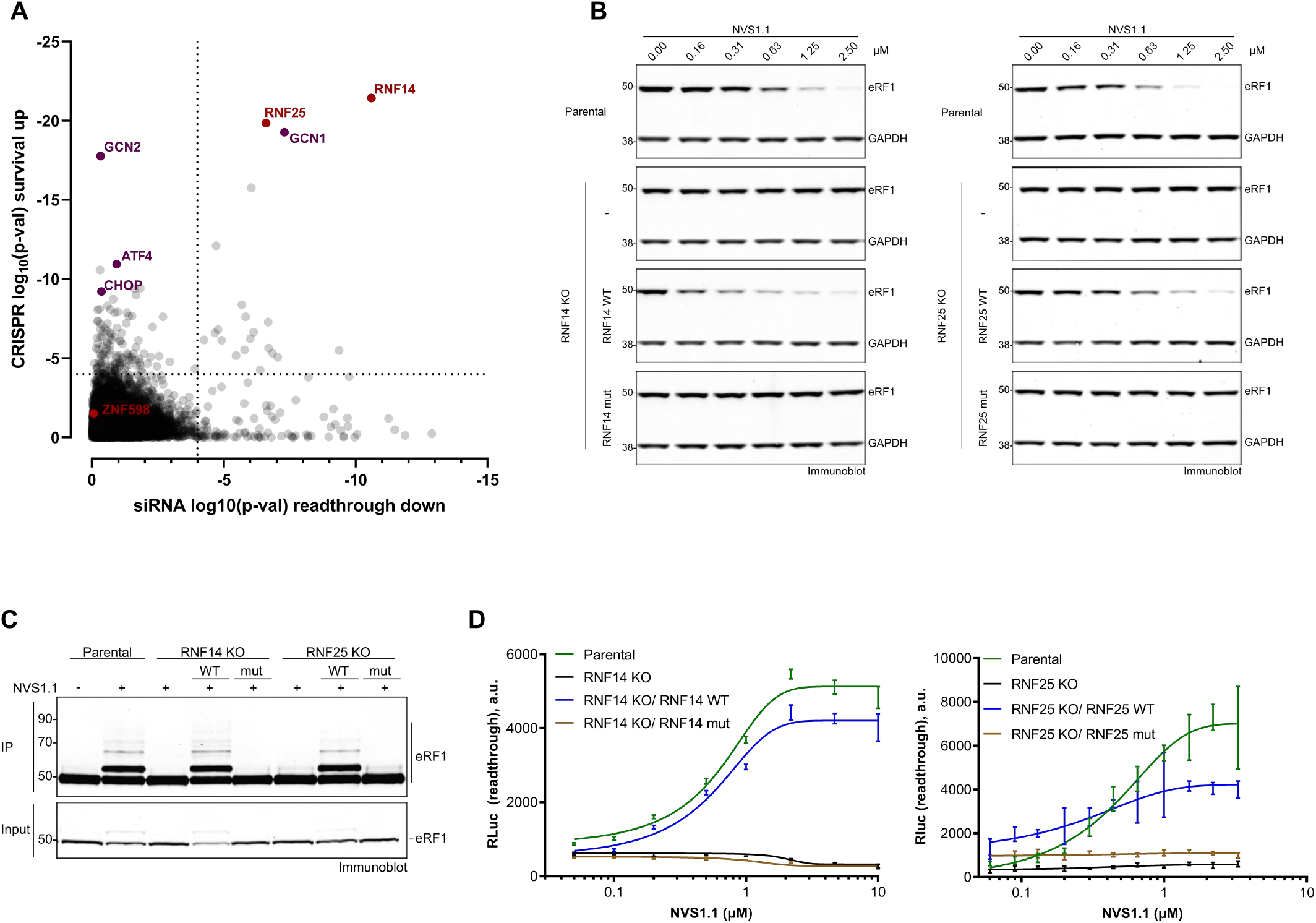
Degradation of eRF1 by NVS1.1 depends on the catalytically active E3 ligases RNF14 and RNF25. (A) Combined results of a genome-wide siRNA screen scoring for reduced NVS1.1-mediated readthrough and a CRISPR knockout screen enriching for genes leading to NVS1.1 resistance. X-axis: Log_10_ p-values depicting the significance of the reversion of NVS1.1-induced readthrough in CFTR-Y122X-Rluc reporter gene expressing HEKR4 PTC cells for the knockdown of each of the 19’300 tested genes. For each gene knockdown (8 siRNAs per gene), Rluc activity was normalized to the luminescence signal of a non-targeting siRNA (log_2_FC) and the differential activity between the two treatment conditions (NVS1.1 IC_80_ vs. DMSO) was determined. The gene significance was calculated for the differential activity of each gene knockdown using the RSA statistical test (redundance siRNA activity) ^67^. Y-axis: Log_10_ p-values depicting the significance of the reversion of NVS1.1-induced toxicity in a cell survival assay for the knockout of each of the 19’300 tested genes. Differential representation of each sgRNA in NVS1.1 IC_80_ and untreated library-infected cell populations was determined as surrogate of difference in cell proliferation. The gene significance was calculated for the differential representation of each sgRNA set (5 sgRNAs per gene) using the RSA statistical test. For both screens, the significance thresholds were determined by randomizing the gene labels before running the RSA tests. A log_10_(p-Val) <-4 threshold (dotted lines) limited false positives to ∼5%. (B) Knockouts of the E3 ligases RNF14 and RNF25 were generated in the HEKR4 PTC reporter cell line (RNF14KO and RNF25KO, respectively). From these knockout cells, rescue cell lines were created that stably overexpress either the wild-type (WT) protein or a catalytically inactive mutant (RNF14mut [C220S] and RNF25mut [C135S/C138S], respectively). Parental, knockout, and rescue cells were incubated with the indicated NVS1.1 concentrations for 6 hours. eRF1 levels in these cells were assessed by immunoblotting, whereby GAPDH was used as a loading control. (C) To assess the ubiquitination status of eRF1, the parental, knockout and rescue cells described in (B) were incubated with 25 μM NVS1.1 for 30 min, followed by eRF1 immunoprecipitation. 50% of the eluates and 1% of input were then analyzed by immunoblotting using anti-eRF1 antibody. (D) Measurement of translational readthrough (represented as arbitrary units, a.u.) in the parental, knockout and rescue cells described in (B) after incubation with a serial dilution of NVS1.1 for 24 hours.

To validate the involvement of RNF14 and RNF25 in NVS1.1-induced proteasomal degradation of eRF1, we generated clonal knockouts of these genes in the HEKR4 PTC reporter cells and tested the sensitivity of eRF1 for NVS1.1-induced degradation in these cells (Figures 5B and S5A). While in the parental cells, eRF1 protein levels were decreased in a dose-dependent manner, the knockout of RNF14 or RNF25 rendered the cells resistant to NVS1.1-induced eRF1 degradation. Complementation analysis showed that the NVS1.1 sensitivity was restored by the expression of recombinant active RNF14 (WT) but not by catalytically inactive RNF14 C220S mutant (mut), in which a cysteine residue of the RING finger domain critical for E3 ligase activity was changed to serine (Figure 5B and S5A) ^40^. Likewise, RNF25 WT but not the catalytically inactive RNF25 mut (C135S/C138S) ^41^ restored NVS1.1 sensitivity of eRF1 in the RNF25 knockout cells (Figure 5B and S5A).

eRF1 ubiquitination was only detected in cells expressing active RNF14 and RNF25 but not in cells depleted for one of the two E3 ligases or in cells expressing mutated versions of RNF14 or RNF25, demonstrating that RNF14 and RNF25 are non-redundant E3 ubiquitin ligases required for ubiquitinating eRF1 in response to NVS1.1 (Figure 5C). That both E3 ubiquitin ligases are required for eRF1 polyubiquitination furthermore excludes a scenario in which one E3 ligase would catalyze the mono-ubiquitination and the other E3 ligase extends ubiquitin chains. In accordance with the requirement of RNF14 and RNF25 for eRF1 degradation, both E3 ligases also proved essential for NVS1.1-induced readthrough. Readthrough of the PTC of the RLuc reporter gene used in the initial screen (Figure 1A) increased in a NVS1.1 dose-dependent manner in parental cells and in cells, in which the knockout of RNF14 or RNF25 was rescued by overexpressing the respective WT proteins. In the RNF14 and RNF25 KO cells overexpressing the respective mutant proteins, only background luminescence was detected, suggesting they are not responsive to the readthrough molecule (Figure 5D). Collectively, our data demonstrates that the extent of NVS1.1-induced translational readthrough is directly dependent on the intracellular concentration of eRF1, which is regulated by RNF14 and RNF25.

Despite being chemically very different from NVS1.1, the mode of action of NVS2.1 appears to be identical (Figure S5A): the siRNA and CRISPR screens conducted with NVS2.1 also identified RNF14 and RNF25, albeit the latter did not pass the threshold filter in the siRNA screen (Figure S5B).

### NVS1.1 triggers a GCN1-mediated ribosome-associated quality control by trapping eRF1 on terminating ribosomes

In addition to the E3 ubiquitin ligases RNF14 and RNF25, the CRISPR knockout screen revealed GCN1 (General Control Non-derepressible 1), GCN2 (also known as EIF2AK4), ATF4 and CHOP as top hits (Figure 5A). GCN1 is a positive regulator of the protein kinase GCN2, which is implicated in the activation of the integrated stress response (ISR), leading to the inhibition of translation initiation by phosphorylation of eIF2-α. This repression of global protein synthesis is accompanied by the specific upregulation of ATF4 and other stress response factors ^42-44^. Depending on the level and duration of ISR activation, different cellular fates can be promoted, ranging from pro-survival signaling aiming at cellular recovery in reaction to short-lived stresses, to programmed cell death upon sustained ISR ^43^. Therefore, it is plausible that during substantial eRF1 depletion, loss of ISR factors increased survival by delaying cell apoptosis conferring an indirect NVS1.1 resistance to these cells in the CRISPR knockout screen. A similar observation has been reported for CC-90009, a molecular glue degrader that leads to the depletion of translation termination factor eRF3A ^34^. However, in contrast to the other identified ISR factors, GCN1 was additionally strongly enriched in the siRNA screen, which scored for readthrough inhibition (Figure 5A). This argues for a direct role of GCN1 in the NVS1.1-triggered eRF1 degradation. Indeed, knockdown of GCN1 in an independent experiment caused a marked stabilization of eRF1 in the presence of NVS1.1 (Figure 6A). Therefore, we conclude that GCN1 is a key component of the pathway by which NVS1.1 causes eRF1 degradation, together with the E3 ligases RNF14 and RNF25.

**Figure 6.**
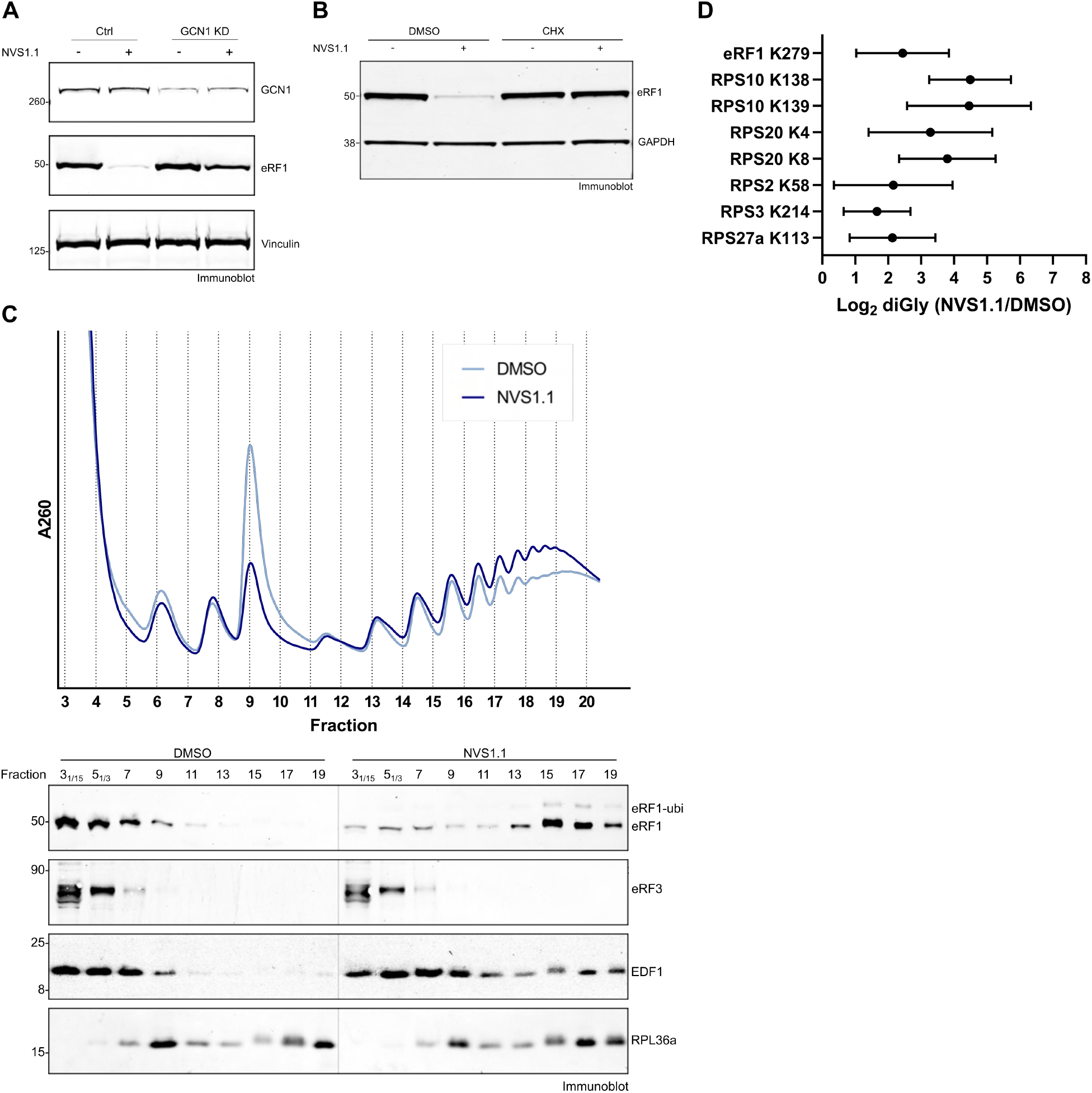
NVS1.1 triggers GCN1-mediated RQC by trapping eRF1 on terminating ribosomes. (A) Immunoblot showing the protein levels of GCN1 and eRF1 in HEKR4 PTC reporter cells treated with negative control (Ctrl) or GCN1-targeting siRNA (GCN1 KD) in combination with a 6-hour incubation of 2.5 μM NVS1.1 or DMSO as control treatment. Vinculin served as a loading control. (B) Representative immunoblot showing protein levels of eRF1 and GAPDH (loading control) of HEKR4 PTC reporter cells treated with 2.5 μM NVS1.1 for 6 hours in the absence or presence of the translation elongation inhibitor cycloheximide (CHX). (C) (Top) Polysome profile showing A_260_ readout of lysate deriving from HEKR4 PTC reporter cells treated with 25 μM NVS1.1 for 30 min that was separated over a 15%-50% sucrose gradient and fractionated. (Bottom) The proteins in every odd-numbered fraction of the gradient were precipitated and analyzed by immunoblotting for the indicated proteins. The fractions 3 and 5 were diluted 1/15 and 1/3, respectively. (D). Heavy polysome fractions from (C) were pooled and the proteins were analyzed by label-free mass spectrometry. Detected diGly events on lysine residues indicative of ubiquitination were normalized to the corresponding protein abundance. The fold change of the diGly frequency (log_2_ diGly (NVS1.1/DMSO)) is shown for the protein eRF1 and for ribosomal proteins that showed a statistically significant difference (pVal < 0.05) upon NVS1.1 treatment

In mammals, the ISR can be activated by multiple kinases upon different stimuli ^43^. Among those kinases, the GCN1-GCN2 branch responds to translational disturbances such as amino acid starvation, UV treatment and alkylating agents, which were shown to induce ribosome collisions ^45,46^. For robust ISR activation, binding of GCN1 to ribosomes is required ^47^. Corroborating these findings, a cryo-EM structure revealed how GCN1 binds to stalled and collided disomes, suggesting that it acts as a collision sensor ^48^. In light of GCN1’s reported connection to ribosomal stalling and collisions, we hypothesized that NVS1.1 could directly influence translation termination and that eRF1 degradation might be a consequence of ribosome stalling at the stop codon. Consistent with this hypothesis, treatment of cells with the translation elongation inhibitor cycloheximide (CHX) rendered eRF1 immune to NVS1.1-mediated degradation (Figure 6B), revealing that the NVS1.1 mode of action relies on actively translating ribosomes. Next, we investigated whether NVS1.1 treatment changes the overall distribution of ribosomes on mRNAs by assessing polysome profiles from NVS1.1 and DMSO-treated cells. These experiments revealed that NVS1.1 reduces the monosome peak and concomitantly increases the heavy polysome fraction (Figure 6C, top panel), potentially resulting from ribosome stalling accompanied by ribosome collisions. Under control conditions (DMSO), eRF1 is predominantly found in the light fractions of the gradient and hardly detectable in the polysome fractions. In contrast, NVS1.1 treatment caused a striking shift of partially ubiquitinated eRF1 into the heavy polysome fractions (Figure 6C, lower panel), suggesting that NVS1.1 traps eRF1 on ribosomes and inhibits translation termination. By contrast, eRF3 was not observed to co-migrate with ribosomes and remained in the light fractions when NVS1.1 was added (Figure 6C, lower panel). Considering that eRF1 and eRF3-GTP mediate translation termination as a ternary complex ^49^, the absence of eRF3 accumulation in the polysome fractions of NVS1.1-treated cells indicates that translation termination in these cells can proceed until GTP hydrolysis and subsequent eRF3 dissociation but is blocked before eRF1 can leave the ribosome. Finally, we assessed the distribution of EDF1, a protein that is recruited to polysomes in a collision-dependent manner and that recruits the translational repressors GIGYF2 and 4EHP to collided ribosomes to prevent new ribosomes from initiating translation on defective mRNAs ^50,51^. In our polysome fractions, EDF1 was mainly detected in the light, ribosome-free fractions and, to a lesser extent, in the monosome fraction under control conditions (DMSO). However, when cells were treated with NVS1.1, a portion of EDF1 migrated deeper into the gradient and was detected in the polysome fractions (Figure 6C, lower panel). Collectively, the accumulation of ubiquitinated eRF1 and EDF1 on polysomes suggests that NVS1.1 inhibits eRF1’s function in translation termination, leading to ribosome stalling at stop codons and subsequent collisions with trailing ribosomes.

Ribosome collisions are known to induce ubiquitination of ribosomal proteins, which serve as important signals for a multi-layered downstream quality control process that mediates ribosome rescue and recycling ^52^. To assess whether NVS1.1-induced ribosomal stalls cause similar ubiquitin signatures, we analyzed the heavy polysome fractions by label-free mass spectrometry. Confirming our immunoblot result, a comparative analysis of the protein content of the two conditions confirmed the accumulation of eRF1 on polysomes when cells were treated with NVS1.1 (Figure S6A). Likewise, the ribosome collision sensor EDF1 was also enriched in the polysome fractions upon NVS1.1 treatment, even though the statistical significance threshold was not reached due to variation between the replicates (Figure S6A). GCN1 was detected at similar levels in the polysome fractions of control and NVS1.1-treated cells, indicating that the GCN1 interaction with ribosomes *per se* is not the activation step for NVS1.1-mediated eRF1 degradation. Furthermore, the relative abundances of ribosomal proteins also remained unchanged, suggesting that treatment with NVS1.1 does not alter the overall ribosome composition.

While the abundance of ribosomal proteins in polysomes was not affected by NVS1.1, inspection of post-translational modification events revealed multiple NVS1.1-specific ubiquitination events (detected as diGly remnants) on small subunit ribosomal proteins, additionally to the previously identified eRF1 K279 ubiquitination (Figure 6D). Among the most enriched modifications, we found ubiquitination of RPS10 (eS10) on amino acids K138 and K139 and of RPS20 (uS10) on K4 and K8. Ubiquitination of RPS10 and RPS20 is mediated by the ubiquitin E3 ligase ZNF598, which is thought to recognize the characteristic interface of collided ribosomes and thereby acts as a sentinel of translation by recognizing ribosome stalls and collisions ^53-55^. To test whether ZNF598 was also required for NVS1.1-mediated eRF1 degradation, we incubated HEK293 cells in which ZNF598 was knocked out with NVS1.1 and observed an unperturbed eRF1 depletion (Figure S6B). Thus, whereas ubiquitination of the ribosomal proteins RPS10 and RPS20 provides further evidence that NVS1.1 causes ribosome collisions, ZNF598 itself is not directly involved in the degradation of eRF1, which is consistent with the results of the CRISPR knockout screens, in which ZNF598 was not a hit (Figures 5A and S5B). Ubiquitination of two other small subunit proteins in our dataset, RPS2 (uS5) and RPS3 (uS3), was previously reported upon treatment of cells with various proteostasis stressors, including translation inhibitors and ISR agonists ^56^. RNF10 was identified as the ubiquitin E3 ligase responsible for the ubiquitination of RPS2 and RPS3 and for regulating 40S subunit turnover in concert with the deubiquitinating enzyme USP10 ^56,57^. The activation signal for the ubiquitination of RPS2 and RPS3 appears to be distinct from ZNF598, hinting at translation initiation defects rather than elongation stalls, but the mechanistic details have yet to be revealed ^56,57^. Finally, we detected increased RPS27A (eS31) ubiquitination on lysine residue K113. Interestingly, RPS27A K107/K113 ubiquitination was previously observed upon treatment of cells with a ribosome collision-inducing low concentration of emetine ^51^. Furthermore, loss of USP16 – a deubiquitinase involved in 40S ribosomal subunit maturation – led to the accumulation of K113-ubiquitinated RPS27A in a translation-dependent manner ^58^. While the E3 ligase responsible for RPS27A ubiquitination remained elusive, our data hints at RNF14 and/or RNF25.

In summary, our results reveal that NVS1.1 arrests translation by trapping eRF1 on terminating ribosomes, leading to eRF1 degradation by a newly discovered pathway involving GCN1 and the E3 ligases RNF14 and RNF25. In parallel, the resulting ribosome collisions and translational repression trigger a regulatory ribosomal ubiquitination program that involves different previously identified ribosome-associated quality control mechanisms.

## Discussion

In this study we set out to develop novel readthrough compounds with favorable pharmacokinetics and test their efficacy in cellular and animal disease models for cystic fibrosis and Hurler syndrome (Figures 1-3). The combination of unbiased high throughput screening using cellular reporter models and compound triaging with multiple target specific functional assays enabled the selection of promising small molecular weight compounds with similar mode of action. The early use of assays that can also be applied for efficacy measurements in patient-derived primary cells and respective animal tissues thereby increased the chance of identifying attractive chemical starting points. Particularly the assessment of the lead compounds in Hurler patient fibroblasts facilitated the selection of promising scaffolds for chemical optimization and for *in vivo* tool compound characterization. In line with this screening strategy, the two most efficacious compounds, NVS1.1 and NV2.1, are structurally diverse, have different pharmacokinetics and chemical properties but act mechanistically very similar (Figures 4-6). Different compound profiles with a similar mode of action are an ideal scenario to investigate the potential of stop codon readthrough compounds in multiple diseases. We concentrated our early *in vivo* pharmacokinetic and pharmacodynamic assessments on using an engineered Hurler rat model that recapitulates the classical Hurler phenotype and biomarkers ^30^. Despite the relatively modest amounts of restored full-length IDUA enzyme, NVS1.1 and NVS2.1 treatment resulted in a pronounced and dose-dependent reduction of total GAG and of enzymatic GUSB activity in brain tissue. To determine the clinical potential of our readthrough promotors, additional efficacy readouts, such as a reduction of the typically enlarged lysosomal structures and the reduction of inflammatory responses will be needed. Furthermore, given that both compounds primarily trigger the depletion of eRF1, a detailed safety assessment of NVS1.1 and NVS2.1 will be a pivotal part of determining their clinical potential.

Our investigation into the mode of action of NVS1.1 uncovered a GCN1-dependent mechanism that senses terminating ribosomes stalled at stop codons upon NVS1.1 treatment and that activates a novel pathway, in which the ubiquitin E3 ligases RNF14 and RNF25 are required to mark eRF1 for proteasomal degradation. Furthermore, we detect hallmarks of previously described RQC and stress response pathways, including ZNF598-mediated ubiquitination of ribosomal proteins, and NVS1.1-dependent EDF1 co-migration with polysomes, implying a multifaced cellular response pathways activated by NVS1.1-mediated inhibition of translation termination.

Interestingly, remarkably similar observations as ours were reported very recently from investigating the cyclic peptide ternatin-4, an allosteric inhibitor of the elongation factor 1α (eEF1A) ^59^. The authors of this study found ternatin-4 to induce the rapid degradation of eEF1A by a pathway that also involves GCN1, RNF14 and RNF25. Paralleling our findings of NVS1.1-induced eRF1 ubiquitination and proteasomal decay, they showed that RNF14 and RNF25 promote eEF1A ubiquitination upon treatment of cells with ternatin-4. By combining the data of their study with our results, we propose the following working model for the mode of action of NVS1.1 (Figure 7A): NVS1.1 traps eRF1 on terminating ribosomes, which causes collisions of trailing ribosomes with the stalled lead ribosome. These collisions are sensed by GCN1, which triggers ubiquitination of eRF1 in a process that requires the cooperative activity of the E3 ligases RNF14 and RNF25. Analogously to how GCN1 interacts with GCN2, both RNF14 and RNF25 feature N-terminal RWD domains that could interact with the RWD binding domain (RWDBD) of GCN1 ^60,61^. Indeed, using reciprocal immunoprecipitations with overexpressed bait proteins, Oltion and colleagues confirmed the binding of RNF14 (but not RNF25) with the RWDBD of GCN1. Since RNF14-GCN1 interaction occurred in the absence of the elongation stall-inducing agent ternatin-4 ^59^, it is likely that the formation of the RNF14-GCN1 complex is not sufficient to induce eRF1 degradation and that an additional cue is needed. To this end, we identified a set of ubiquitinated ribosomal proteins in response to the NVS1.1 treatment. Some of them are known markers for ribosome-associated quality control events (RPS2, RPS3, RPS10, RPS20), while the role of RPS27A K113 ubiquitination is less well understood. Interestingly, ubiquitination of RSP27A at K107 and K113 was also detected in ternatin-4 treated cells. This ubiquitination was dependent on RNF25 ^59^, which corroborates the notion that ubiquitination of ribosomal proteins is an integral part of the signaling leading to the degradation of eEF1A and eRF1. Notably, the cryo-EM structure of GCN1 bound to two collided ribosomes found the RWDBD of GCN1 to locate near the ubiquitination sites of RPS27A on the small ribosomal subunit ^48,59^ (Figure 7B), suggesting that ubiquitinated RPS27A could function as an allosteric activator of the GCN1-RNF14 complex. Collectively, our data indicates that the eRF1 ubiquitination in response to NVS1.1 requires multiple molecular inputs, including recognizing ribosome collisions by GCN1 and ubiquitination of ribosomal proteins – possibly mediated by RNF25 on RPS27A – as activating signal for RNF14-mediated eRF1 ubiquitination. Although further biochemical investigation is necessary to dissect the mechanistic details of these events, our model provides a rationale for why two E3 ligases are required to degrade eRF1. Furthermore, this multistep activation might represent a mechanistic barrier that prevents induction of eRF1 and eEF1A degradation during stochastically slow but functional termination and elongation events, respectively.

**Figure 7.**
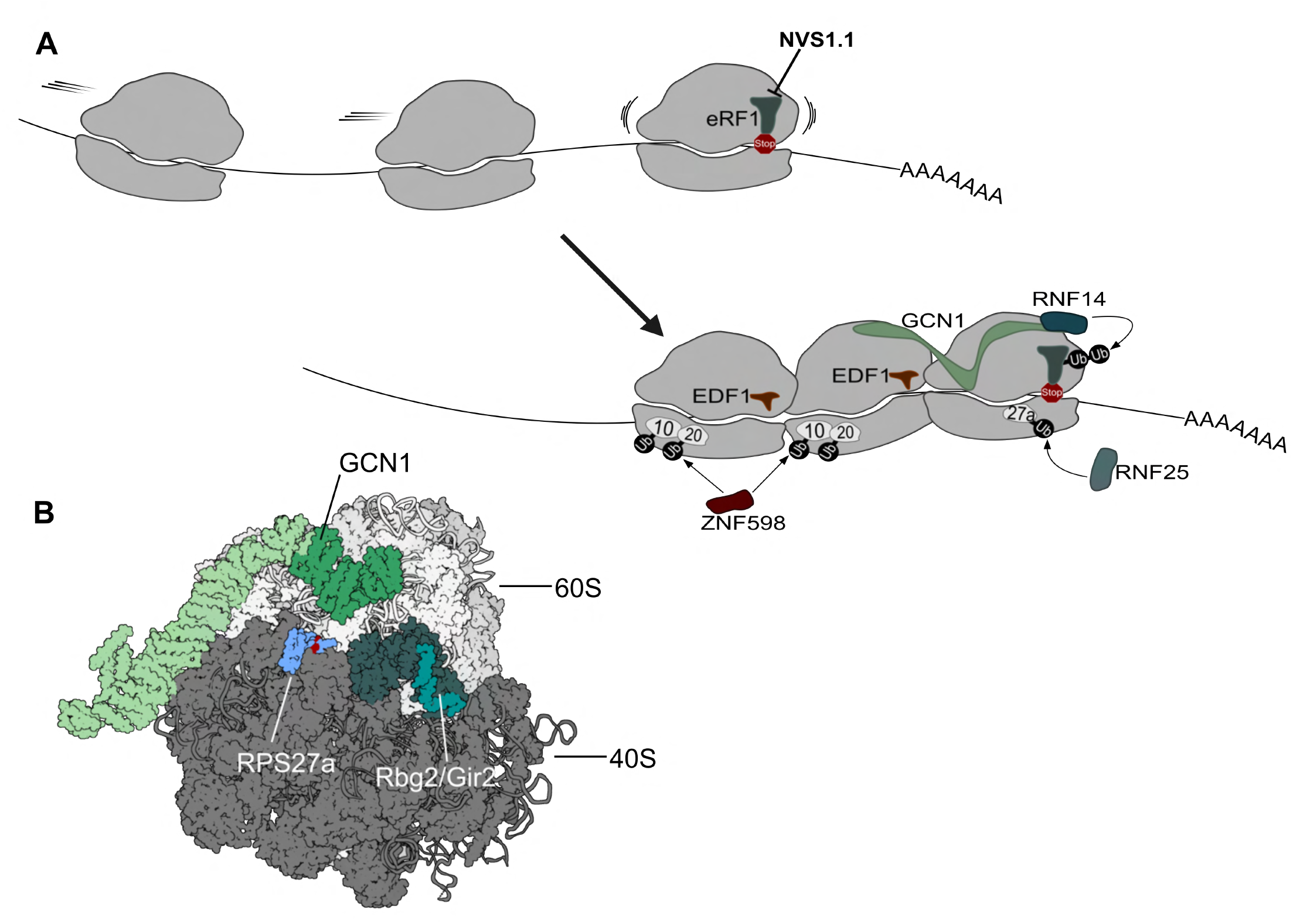
Model for the NVS1.1 mode of action. (A) Schematic model illustrating the GCN1-RNF14-RNF25-dependent nexus of translation quality control that resolves ribosomal NVS compound-induced termination stalls by proteasomal degradation of eRF1. (B) Cryo-EM structure of yeast GCN1-bound lead ribosome within a collided disome complex (PDB 7NRC) showing the 40S in dark grey and the 60S subunit in light grey. The lead ribosome spanning portion of GCN1 is depicted in green with its RWDBD domain shaded in dark green. The GTPase Gir2 (teal) in complex with Rbg2 (dark teal) locates to the A site region. The small ribosomal subunit protein RPS27a is depicted in blue with the lysine residues K107 and K113 marked in red.

Despite the commonalities of the factors involved in the responses to ternatin-4 ^59^ and NVS1.1 (our study), they target different proteins for degradation at distinct translation steps, which raises the question of the underlying principle of these two processes. Ternatin-4 inhibits translation elongation by binding to the ternary complex consisting of aminoacyl-tRNA, eEF1A and GTP and so perturbing tRNA accommodation. This traps eEF1A in the ribosomal A site and results in stalled ribosomes ^62^. The binding site of NVS1.1 is currently unknown, yet our results indicate that it blocks translation termination by preventing eRF1 from leaving the ribosome. Thus, while both compounds target translational GTPase complexes, ternatin-4 targets the GTPase eEF1A for degradation, whereas NVS1.1 targets the decoding factor eRF1 and leaves the GTPase eRF3 unaffected. Therefore, we hypothesize that the degradation pathway involving GCN1, RNF14 and RNF25 recognizes and resolves generally stalled (and collided) ribosomes with an occluded A site rather than recognizing specific proteins. This would be mechanistically distinct from previously described pathways that solve translational problems. For instance, the Pelota-Hbs1L complex preferentially acts on stalled ribosomes devoid of mRNA in the A site ^49,63^, and structural data of collided ribosomes that are ubiquitinated by ZNF598 suggest that the primary recognition motif of ZNF598 is the contact of specific ribosomal proteins at the 40S-40S disome interface, independent of the A site occupancy ^54,64^. In conclusion, we postulate that GCN1, RNF14, and RNF25 comprise a new branch of ribosome-associated quality control that resolves ribosome problems arising from an occluded A site during translation elongation and termination.

GCN1 was so far best known for its role in GCN2-dependent activation of the integrated stress response (ISR) upon sensing ribosome collisions ^45,46^. Our findings suggest a broader role of GCN1 as a more general sensor of ribosome stalling that leads to the activation of distinct downstream pathways depending on the exact cause of the stalling. Whereas empty A sites reduce translation initiation via ISR activation, occluded A sites activate the proteasomal degradation of the obstructing protein. In the case of the NVS1.1-mediated inhibition of translation termination, the A site obstructor is eRF1 and its rapid RNF14-RNF25-proteasome-mediated depletion results in increased readthrough of termination codons, explaining why NVS1.1 is a potent readthrough promoter. Additional work is needed to elucidate the links between GCN1- and ZNF598-mediated responses to ribosome stalls and collisions, and the conditions that trigger these pathways. Furthermore, it remains to be investigated how the ribosomal subunits are recycled after eRF1 ubiquitination and at which stage eRF1 is subjected to proteasomal degradation. Apart from their therapeutic potential, the compounds NVS1.1 and NVS2.1 will provide useful tools to further dissect the intricate network of ribosome-associated quality control mechanisms.

## Limitations of the study

While our study demonstrated how the small molecule compounds NVS1.1. and NVS2.1 lead to ribosome stalling at a PTC and to the GCN1-mediated activation of a novel translation quality control pathway, it remains to be elucidated under which physiological conditions occluded A sites occur that trigger this pathway. However, the finding that overexpression of Itt1p, the RNF14 homolog in *S. cerevisiae*, also causes eRF1 depletion ^65^ suggests an evolutionary conservation and physiological functions of this GCN1-RNF14-RNF25-dependent translation quality control.

## Author contributions

JR and NS were the project leads, supervised and coordinated the drug discovery in vitro, in vivo and Novartis MoA work; IS, YI, OG conducted the HTS screens; NS performed the chemical optimizations of the compounds; JS, CG supported cell biology work; CP and IS did the CRISPR screen; XM supported the in vivo and rat cell work; FB was responsible for MS analysis; PC guided safety related work; FS and MS guided the siRNA, SILAC and MoA proteomic experiments. OM supervised the Bern mode-of-action (MoA) studies; LAG performed and analyzed the MoA experiments with contributions from JZ; ACU analyzed the proteomics data; LAG prepared the figures; LAG, OM and JR wrote the manuscript.

## Acknowledgements

The authors are grateful to Jane Nagel at NIBR Cambridge and Sophie Braga Lagache, Natasha Buchs, and Manfred Heller from the PMSCF of the University of Bern for their excellent mass spectrometry services, to the former team and Novartis colleagues Lloyd Klickstein, Ken Huttner, Natalie Dales, Florian Fuchs and Joseph Kelleher for translational medicine, chemistry, and MoA work, to Steffi Harlfinger for guiding the PK/PD in vivo work, and to Evangelos Karousis for his valuable comments to the manuscript. We further acknowledge the ChemG and target ID proteomic groups at Novartis Cambridge, US, and Basel, CH, and the Novartis CBT and DAx management for supporting the project.

The work conducted at the University of Bern was supported by the National Center of Competence in Research (NCCR) on RNA & Disease funded by the Swiss National Science Foundation (SNSF; grant 51NF40-141735, 182880, and 205601), by SNSF grants 310030B-182831 and 310030-204161, and by the canton of Bern (University intramural funding to O.M.).

## Methods

### Experimental Model and Subject Details

#### Cell lines and culture conditions

The parental Jump-In™ HEK293 (Thermo Fisher Scientific, hereafter termed HEKR4) and all its derivatives were grown in medium containing DMEM (Invitrogen, Cat# 41966052), 10% dialyzed FCS (Bioconcept, Cat# 2-01F16-I), 25 mM HEPES (Invitrogen, Cat# 15630056), 1x MEM Non-Essential Amino Acids Solution (Invitrogen, Cat# 11140035), 100 units/ml of penicillin and 100 µg/ml of streptomycin (Invitrogen, Cat# 15140122), and 10 µg/ml Blasticidin S (Invitrogen, Cat# R21001). Primary patient fibroblasts homozygous for the IDUA W402X mutation were obtained from Coriell (GM00798 [female]; Deficient alpha-L-iduronidase; Hurler syndrome; homozygous for a TGG>TAG change at nucleotide 1293 in exon 9 of the IDUA gene [Trp402Ter (W402X)]) and fibroblasts homozygous for the IDUA Q70X mutation were provided by the Telethon foundation (Request-ID #834). These cells were cultured in medium containing MEM (NEAA, no glutamine, Gibco, Cat# 10370021) supplemented with, 2 mM L-Glutamine (Gibco, Cat# 25030149), 10% FCS (BioConcept, Cat# 2-01F16-I) and 100 units/ml of penicillin and 100 µg/ml of streptomycin (Invitrogen, Cat# 15140122). Rat primary fibroblasts (REFs) were grown in DMEM (Gibco, Cat# 11965092) containing 10% FCS, 100 units/ml of penicillin and 100 µg/ml of streptomycin (Invitrogen, Cat# 15140122). All cells were cultured at 37°C in humidified incubators with 5% CO_2._ The NVS compounds, Bortezomib (BTZ, Merck, Cat# 504314), MLN4924 (Selleckchem, Cat# S7109) and Cycloheximide (CHX, Focus Biomolecules, Cat# 10-117) were all dissolved in DMSO, which also served as vehicle control. For long-term treatment up to 7 weeks in the patient and rat primary fibroblasts, the NVS compounds were reapplied every 3^rd^ day with the medium change. Paromomycin (Sigma Aldrich, Cat# P9297) was dissolved in PBS.

#### Engineering of recombinant PTC reporter, CFTR and IDUA cell models

Introduction of recombinant genes into the parental HEKR4 cell line was performed according to the manufacturer’s guidelines (Thermo Fisher Scientific). In brief, HEKR4 contains a single R4 *att*P retargeting sequence and a promoterless Blasticidin resistance cassette (Bsd^R^). To introduce a gene of interest (GOI) into the recombination site, the GOI was cloned into the pJTI-R4-DEST CMV-pA vector and co-transfected with the pJTI™ R4 Int vector for the expression of the integrase. 5×10^6^ cells per well of a 6-well plate were transfected with a mix of plasmid DNA, Lipofectamine™ LTX Reagent with PLUS™ Reagent (Invitrogen, Cat# 15338100) in Opti-MEM (Gibco, Cat# 31985062) according to the manufacturer’s instructions. On the next day, the cells were transferred to T80 flasks and one day later, the growth medium was supplemented with 10 µg/ml Blasticidin S (Invitrogen, Cat# R21001) to select for site-specific integration events (positioning the EF1a promoter sequence in front of Bsd^R^). After three weeks of culturing and Blasticidin selection, single cells were sorted into 96-well plates. Using the Cellavista brightfield Imager system (Syntec) for screening, single clones were identified and proliferated to the T80 format. While maintaining the Blasticidin selection for cell line propagation, the antibiotic was withdrawn from the medium 1-2 days prior to experimental procedures. For each of the integrated constructs, results from one cell clone are shown.

#### Engineering of knockout and rescue cell lines

Candidate E3 ligases were knocked out using CRISPR-Cas9-mediated genome editing methods. Frist, the guide RNA sequences targeting exon 6 of RNF14 and exon 3 of RNF25 were inserted into the pU6-gRNA-SpyCas9-2A-Puro-eGFP plasmid using annealed oligonucleotides (see Key Resource Table) that were ligated into the Esp3I restriction sites. The resulting plasmid was used to transfect HEKR4 PTC reporter cells in 6-well plates with Lipofectamine 2000 (Invitrogen, Cat# 11668019) and PLUS™ Reagent (Invitrogen, Cat# 11514015) in Opti-MEM (Gibco, Cat# 31985062). After 24 h, the medium was supplemented with 2 µg/ml Puromycin (Sigma, Cat#P9620) to select for transfected Cas9 expressing cells (Cas9 is encoded as fusion protein with an eGFP-Puro^R^ cassette intermitted by a P2A skipping sequence). Puromycin selection was maintained for 3 days and subsequently the cells were reseeded as single cell dilutions into 96-well plates. Clonal cell lines derived from single cells were then assessed for the loss of RNF14 and RNF25 expression by immunoblotting.

To reintroduce the E3 ligases in the background of the knockout cell lines, the open reading frames of RNF14 and RNF25 were PCR amplified and cloned into the pcDNA3.1(+) vector. For both E3 ligases, enzymatically inactive RING domain mutants were created (RNF14 C220S ^40^ and RNF25 C135S/C138S ^41^). The resulting plasmids were transfected into RNF14 KO cells or RNF25 KO cells using Lipofectamine 2000 (Invitrogen, Cat# 11668019) and PLUS™ Reagent (Invitrogen, Cat# 11514015) in Opti-MEM (Gibco, Cat# 31985062). On the next day the growth medium was supplemented with 400 µg/ml Geneticin (G418, Thermo Fisher Scientific, Cat# BP673-5) to select for random genomic incorporation of the plasmids. Selection was maintained for about three weeks, before single cell-derived clones were raised for the RNF14 WT/mutant rescue cell lines as described above and assessed by immunoblotting. For the RNF25 WT/mutant rescue cell lines, protein overexpression was verified by immunoblot in the cell pools, which were then directly used for experiments. Blasticidin and G418 selection was maintained during cell line propagation and withdrawn from the medium 1-2 days prior to experimental procedures. Rluc readthrough measurements with RNF14, RNF25 KO and overexpressing cells was done as described in the HTS method section.

#### siRNA-mediated knockdown of GCN1

Approx. 4×10^6^ HEKR4 PTC reporter cells were grown on a 6-well plate and transfected with 40 nM of negative control (NegC, siPool) or GCN1 (siPool, 10985 – GCN1, human) targeting siRNA in a 200 μl serum-free mix containing 12 μl Lullaby (Oz Biosciences, Cat# LL71000), resulting in a final volume of 2 ml after addition to the cells. After one day, each condition was split 1:4 and one day later the transfection was repeated. One day after the second transfection the cells were treated with DMSO or 2.5 μM NVS1.1 for 6 h and immediately harvested by scraping in cold 1x PBS. The cells were then collected by centrifugation at 500 x *g* for 5 min at 4°C and stored at -80°C until further processing.

#### High throughput screening and hit filtering

Screening (1.6 million compounds, 10 µM) was performed with the described HEKR4 GFP-CFTR-Y122X-hRluc cell line. Cell culture was supported by a fully automated cell culture device (SelecT, TAP_UK) and screening was done in 1536 well plate assay format (Greiner, Cat. #789183-A). Cell passaging was done every 3^rd^. day and cell were used between cell passage 9-22. *Renilla* luciferase (RLuc) readout was carried out in a fully automated uHTS screening factory equipped with Envision (Perklin Elmer) readers. Cells were dispensed by the SelecT device (2’000 cells/well in 4 µl) and incubated for 24 h (37°C, 5% CO_2_). Compounds were applied at a final concentration of 10 µM using an automated uHTS PinTool device (dispensing 40 nl compound solution per well). Control compound (Paromomycin 14.4 mM f.c) was dispensed with a Flexdrop device to plate column 47 and 48. *Renilla-*Glo assay substrate (Promega, Cat. #E2750) was dispensed using an automated and customized uHTS Multidrop device (2.5 µl/well). Plates were slightly centrifuged at room temperature for 2 min on the screening factory. RLuc signal as readout for compound mediated translational readthrough was recorded with the Envision reader (0.1 s, US-Lum protocol, 0.1 mm distance aperture). Confirmation (20 µM, n=4) and validation screening (n=3, 8-point dose-response) was done offline using the very same readers. Offline compound transfer was done with an acoustic Echo dispenser (10-40 nl). Screening analysis and hit list selection was done with Novartis proprietary software. Z’ factor calculation was done with the formular Z’ = 1-3* stdv High value) + 3* stdv (Low)/Average (High) – Average (Low) as previously described ^66^. Screening hits were identified with the formular: A1 (%) = 100*(S-NC)/ (AC-NC) where AC, NC and S correspond to Active Controls (injection of Stimulation buffer = 100% stimulation), Neutral controls (buffer injection EC_10_) and screening samples (S). NC corresponds to 0% activity whereas AC is 100% activity (full stimulation). Post hit confirmation, screening hits were filtered with a HEKR4 GFP-CFTR-WT-Rluc cell line.

The compounds were triaged with an immunofluorescence imaging high content assay monitoring the restoration of the PTC mutant coagulation factor 9 (F9) R338X, stably expressed in HEKR4 cells. A corresponding F9-WT expressing HEKR4 cell line was used as reference. For AC (active control = 100%) control NVS2 was used. As screening neutral control (NC) DMSO solution (0.5% in media) was applied. Cell culture medium for F9 filtering was identical to the HTS Rluc assay. Compound transfer from compound source plates was done with an Echo device into 1536 well black clear bottom plates (BD Cat. #359315) containing 2’000 cells/well in 6 µl volume (cell media). Final compound concentration for confirmation screening was 10 µM (n=4). For concentration response the compounds were diluted in 90% DMSO as 8-step dilution series and a dose range between 50 µM and 36 nM. For detection of F9 restoration paraformaldehyde-fixed (15 min, 4.4 % f.c. in PBS) cells were washed twice with PBS, incubated with permeabilization and blocking solution (20% FCS, 0.2% Triton X-100 in PBS) for 30 min at room temperature and incubated with an anti-F9 antibody (1:600 in PBS, 1% FCS) together with a DRAQ5 (1:4’000) nuclear stain for 1 h at room temperature. Cells were washed with PBS and an anti-mouse IgG AF488 (1:300 in PBS, 1 % FCS). Cells were washed twice with PBS and imaged (Opera imager) using dual excitation at 647 nm and 488 nm wavelength and emission at 690 nm and 549 nm for DARQ5 and antibody staining, respectively. IF data was analyzed with a proprietary imaging software. F9-containing cytoplasmic regions of interest were framed and nuclei imaging was excluded (% Factor IX+ cells based passing the cytoplasmic intensity threshold). Z’ factor of the assay was > 0.5.

#### siRNA screen

HEKR4 PTC reporter cells grown as described above were used for the siRNA screen. The siRNA library, designed to target 19’300 genes whereby 8 different siRNAs were used per gene. siRNAs dissolved in Opti-MEM (Gibco, Cat# 31985062) were dispensed into 384 well plates, 75 nl/well using the Echo Series Acoustic Liquid Handler (Beckman) at room temperature. Lipofectamine™ RNAiMAX Transfection Reagent (Invitrogen, Cat# 13778075) was diluted 1:266 in Opti-MEM and 2 μl of this mixture was added to each well. Then 2 μl cell suspension containing 10’000 cells was transferred to each well. Finally, 10’000 cells (2 μl) were seeded into each transfection mix-containing well and incubated at 37°C, 5% CO_2_ in humid atmosphere. One day later compound treatment was started by addition of NVS1.1 or NVS2.1 (3 μl/well) resulting in final concentrations of 1.21 μM or 6 μM, respectively which corresponds IC_80_ of each compound to the previously determined in the recombinant cell models. DMSO was used as vehicle control. After 48 h of compound incubation, the luciferase expression was assessed (Promega, Cat# E2710) by addition of 3 μl Renilla-Glo™ reagent (Promega, Cat# E2710) and measuring the luminescence signal with the ViewLux uHTS Microplate Imager (PerkinElmer, Cat# 1430-0010A). The raw data was analyzed by a Novartis proprietary screening software as follows: Activity calculated as siRNA activity (Rluc) divided by median activity of plate negative control siRNA wells, Robust z-score normalization on each plate (robust z-score = (log2_FC – Median_Log2_FC) / (MAD_Log2_FC*1.4826). Statistical tests were performed using the RSA-analysis (redundance siRNA activity) ^67^.

#### CRISPR screening procedures

CRISPR screen was done in GFP-CFTR-Y122X-Rluc reporter cells. Cas9 was stably expressed using the plasmid pLenti6P-CMV-3xFLAG-NLS-SPyCas9_NLS-t2a-Hygro. Stable cell clones were derived after 2 weeks of Hygromycin selection. Cas9 expression was confirmed by immunofluorescence with paraformaldehyde (4% f.c, 15 min, room temperature, Electron Microscopy Sciences, Cat. #15713-S) fixed cells. Blocking and antibody dilution was done in 3% bovine serum albumin (BSA) solution in DPBS (Sigma-Aldrich #A7979), 0.1% Triton X-100 (Sigma-Aldrich, Cat# 93443) solution in H_2_O. The Cas9 protein contained an N-terminal FLAG-tag to confirm expression (mouse anti-3x FLAG primary antibody, 1:250 dilution, RT, 3 h, Sigma-Aldrich Cat# F1804). Cells were washed with PBS buffer. The anti-FLAG antibody was detected using an anti-mouse Alexa 647 secondary antibody (Alexa 488 goat anti-mouse, 1:500 dilution, RT, 1 h, Invitrogen, #A11029). The images were captured with an InCell2000 imager.

##### Cell Line sensitivity to NVS1.1 and NVS2.1

Cells, 5×10^5^, were seeded into a 6 well format the day before compound addition. NVS1.1 was tested at 0, 1.5, 3, 6, 8 and 10 µM and NVS2.1 at 0, 6, 12, 24, 30 and 35 µM. The cells were incubated for 8 days with two changes of media containing the appropriate compound concentration. Cell growth and death was monitored using microscopy imaging using GFP and bright field (BioRad ZOE Imager) and Cell TiterGlo. The final compound screening concentration was chosen at 3 µM for NVS1.1 and 24 µM for NVS2.1 (i.e., between 10-20-fold greater than the Reporter-RLuc IC_50_ of each compound).

##### Library Transduction

For each compound screening two cell stacks (Corning, Cat# 3319) each containing 6.7×10^7^ of cells were seeded the day before virus infection. At the day of the infection, 700 ml cell culture medium per cell stack was completed with polybrene (8 µg/ml f.c.), mixed, incubated for 5 min before adding the calculated amount of virus stock (7 ml for C-pool 1 CP1004, 8 ml for C-pool 3 CP3001). The cell culture medium of the cell stacks was removed, and the virus-containing medium was poured into the cell stacks. The cell stacks were incubated at 37°C, 5% CO_2_ for 24 h before exchanging the medium and adding puromycin-containing medium (4 µg/ml, final concentration). The transduction efficiency was monitored for five days post infection by RFP-positive cells measured by FACS Aria. Viral pools achieved > 92% expression.

##### Preparation of Genomic DNA

Genomic DNA was prepared using Qiagen’s QIAamp DNA blood maxi kit (Qiagen Cat #51194) as recommended by the manufacturer. The genomic DNA was eluted in 1 ml of buffer AE. To ensure complete recovery of the DNA, an extra 1 ml of buffer AE was added to the column and recovered again by centrifugation for 5 min.

##### DNA quantification

Genomic DNA (gDNA) was quantified using the Quant-iT PicoGreen dsDNA assay (Invitrogen) following the manufacturer’s recommendations. In short, the lambda DNA standard is diluted to a working concentration of 2 µg/ml using TE buffer. In an optically clear flat bottom 96 well plate a 10 point 100 µl volume lambda DNA standard curve is created by serially diluting the lambda DNA working solution using TE buffer. The gDNA is diluted in 100 µl TE buffer in an optically clear flat bottom 96 well plate. A working solution of PicoGreen is created by diluting the reagent 200-fold using TE buffer. 100 µl of the PicoGreen working solution is added to the lambda DNA standards and the diluted gDNA samples. The samples are mixed, incubated for 2-5 min at room temperature, protected from light and the fluorescence is measured using a fluorescence microplate reader (Envision, Perkin Elmer) and standard fluorescein wavelengths following the manufacturer’s recommendations. The gDNA sample concentration is then determined using the lambda DNA standard curve.

##### Illumina library construction

To determine the gRNA representation of the lentiviral gRNA transduced samples (and the input gRNA plasmid library), the integrated gRNA sequences were PCR-amplified and sequenced using the Illumina sequencing technology. Illumina sequencing libraries are generated using PCR with primers specific to the integrated lentiviral vector sequence. PCR primers also contain additional sequences required for Illumina sequencing and sample multiplexing. It was empirically determined that a total of 96 µg of gDNA (an average of ∼300 cells per gRNA), divided into 24, 4 µg PCR reactions, is required to accurately determined the representation of 55,000 gRNA sequences within a sample. PCR reactions are performed in a volume of 100 µl, containing a final concentration of 0.5 µM of each PCR primer (Integrated DNA Technologies, 5644 5’-AATGATACGGCGACCACCGAGATCTACACTCGATTTCTTGGCTTTATATATCTTGTGGAAAGGA-3’ and INDEX 5’-CAAGCAGAAGACGGCATACGAGATXXXXXXXXXXGTGACTGGAGTTCAGACGTGTGCTCTTCCGATC

-3’, where the X denote a 10 base PCR-sample specific barcode used for data demultiplexing following sequencing), 0.5 mM dNTPs, 1x Titanium Taq DNA polymerase and buffer (Clontech). PCR cycling conditions used are: 1x 98°C for 5 min; 28x 95°C for 15 sec, 65°C for 15 sec, 72°C for 30 sec; 1x 72°C for 7 min; and a final 4°C hold. The resulting Illumina libraries are purified using 1.8x SPRI AMPure XL beads (Beckman Coulter) following the manufacturer’s recommendations. In short, 5 µl from eight independent PCR reactions are pooled together, generating three 40 µl pools (each pool containing eight PCR reactions). Each 40 µl pool is combined with 72 µl (1.8x) SPRI AMPure XL beads, mixed and incubated for 5 min at room temperature. Samples are then placed on a magnet (tube or plate) and left for 2 min at room temperature. The supernatant is carefully removed without dislodging the beads. The beads are washed with 200 µl fresh 75% ethanol for 30 sec and the supernatant is carefully removed without dislodging the beads. The beads are air dried to remove any excess ethanol for 1-2 min at room temperature. Once dry, the samples are removed from the magnet, 50 µl of nuclease water is added, and the beads are mixed and incubated for 2 min at room temperature to elute the Illumina libraries from the beads. The samples are placed back on the magnet for 2 min and the supernatant is carefully removed into a new plate/tube without dislodging the beads.

##### Illumina library quantification and pooling

The library pools are quantified using a SYBR green qPCR with primers specific to the Illumina sequences (Integrated DNA Technologies, P5 5’-AATGATACGGCGACCACCGAGA -3’ and P7 5’-CAAGCAGAAGACGGCATACGA -3’). In short, 7.8 µl of Power SYBR green master mix (Thermo-Fisher), containing a final concentration of 2 µM of each primer (P5 and P7) is added to 7.2 µl of each library pool and subjected to qPCR; this is performed in duplicate. Additionally, a library of known concentration is run alongside as a control. qPCR cycling conditions used are: 1x 95°C for 10 min; 30x 95°C for 15 sec, 60°C for 1 min. For the qPCR data analysis, a baseline of 2-8 C_T_ and a threshold of 0.2 is used. C_T_ values from technical duplicates are averaged and converted to a linear C_T_ (LCT) value using the following equation LCT= Exp (average C_T_ - 22.0) / -1.55). The library pools are then further pooled such that a total of four gDNA samples (i.e., 12 pools for 4 gDNA) are pooled for sequencing on one lane of an eight-lane high output HiSeq2500 sequencing flow cell. Pooling is done by pooling together a fixed LCT from each library pool. Each gDNA sample receives 50-60M reads which is equivalent to a ∼1000 reads per sgRNA.

##### Illumina flowcell generation and sequencing

For sequencing a single read HiSeq sequencing flowcell is prepared on the cBot instrument (Illumina), using a TruSeq SR Cluster v4 cBot kit (Illumina) and the ‘SR_AMP_LIN_BLOCK_StripHyb.v9’ cBot program, following the manufacturer’s recommendations with the following changes. Library pools are denatured by combining 17 µl of 10mM TRIS buffer (Qiagen buffer EB) with 1 µl 2M NaOH and 2 µl of the 2000 LCT/ µl library pools and incubated at RT for 5min. A 2 µl aliquot of the denatured library pool is then added to 1 ml of HT1 (Illumina), mixed and 120 µl is added to the ‘sample library’ cBot strip tube. As these libraries require a custom sequencing primer, reagent strip tube #2 in the cBOT reagent kit is replaced with a strip tube containing 350 µl per tube of a custom read 1 sequencing primer (Integrated DNA Technologies, 5645 5’-TCGATTTCTTGGCTTTATATATCTTGTGGAAAGGACGAAACACCG-3’) at a concentration of 0.5 µM (e.g., 15 µl 100 µM 5645 primer in 3 ml HT1). The sample barcode is sequenced using the standard Illumina indexing primer (5’-GATCGGAAGAGCACACGTCTGAACTCCAGTCAC-3’), following the manufacturer’s recommendations. The clustered flowcell is then sequenced with a 30b read 1 and an 11b index read using a HiSeq SBS v4 50 cycle kit (Illumina) following the manufacturer’s recommendations.

##### CRISPR Screening

Due to toxicity of the lead compounds in this cell line it was possible to run this screen as a survival assay looking for genes that, when knocked-out by CRISPR-Cas9 indels, caused either increased sensitivity to the compounds (and hence causing these genes to be further under-represented in the pool of survivors) or alternatively to be lead to resistance to the compounds and hence to be overrepresented in the pool of survivors. Live cells were collected after treatment and gDNA prepared for analysis of those genes over and underrepresented in the surviving pool.

##### Hit analysis

After selection of cells surviving compound treatment, genomic DNA was prepared from the surviving cells and then used for sequencing and analysis of the barcoded CRISPR knockouts. Statistical tests were performed using the RSA-analysis (redundance siRNA activity) ^67^.

#### CFTR expression assessment by immunofluorescence microscopy

HEKR4 cells stably expressing S-tagged wild type CFTR or the PTC mutants Y122X, G542X, W1282X were seeded in lysine coated 384-well plates (Corning, Cat# 359332), 8’000 cells/well. After 6 h, the cells were incubated with serial dilutions of NVS1 (0-25 µM), NVS2 (0-50 µM) or Paromomycin (0-10 mM). DMSO served as vehicle control. After 48 h, the anti-S tag antibody (Merck, Cat# 71549-3) was added directly to the medium of each well resulting in a final dilution of 1:500, followed by 20 min incubation at room temperature. Subsequently, the cells were fixed with 4% paraformaldehyde for 15 min at room temperature, followed by three washes with PBS. The primary antibody was then detected by an Alexa Fluor™ 488-conjugated goat anti-mouse secondary antibody (Invitrogen, Cat# A-11001), which was diluted 1:500 in blocking solution (PBS containing 1% FCS [Bioconcept, Cat# 2-01F16-I]) and incubated at room temperature for 1 h. DRAQ5 (Thermo Fisher Scientific, Cat# 62251) 1:5’000 diluted in 1% FCS was included in the same step to stain the nuclei of the cells. Cells were visualized with 20-63x magnification using the Zeiss LSMII confocal microscope.

#### CFTR membrane potential assay

4’000 HEKR4 cells per well expressing the above-described CFTR variants were seeded into lysine coated 384 well clear bottom plates (Thermo Fisher Scientific, Cat# 142761). 6 h later, 10 µM NVS1 or NVS2, DMSO, or 10 mM Paromomycin was added (final concentrations), and cells were further incubated for 48 h in culture medium without antibiotics. Cells were then washed with a Na-Gluconate loading buffer containing 120 mM Na-Gluconate, 2 mM CaCl_2_, (Merck, Cat# 2382), 2 mM MgCl_2_ (Fluka, Cat# 63063), 10 mM HEPES, pH 7.4 at room temperature. The membrane potential dye was diluted as recommended by the manufacturer (Molecular Device, Cat# R8034) in loading buffer. After cell washing, 20 µl membrane potential dye in loading buffer was incubated in each well for 30 min at 37°C. 20 µM Forskolin (Sigma Aldrich, Cat# F6886, dissolved in ethanol) was injected online on a FLIPR (MD) high throughput cellular screening platform. Fluorescence increase was plotted as ΔF/Fb and normalized to the respective WT CFTR response. As a negative control, cells were preincubated for 20 min with 10 µM of the CFTR inhibitor Inh172 (Sigma-Aldrich, Cat# C2292) before Forskolin stimulation.

#### RT-qPCR in hurler patient fibroblasts

Fibroblasts from a female Hurler syndrome patient (IDUA-W402X, Coriell Institute, Cat# GM00798) were grown in 6-well plates to a density of approx. 6×10^5^ cells per well. The cells were then incubated with different concentrations of NVS1.1 ranging from 0.3 μM to 5.0 μM for 8, 16, or 24 h. Thereafter, the cells were collected by trypsinization (Gibco, Cat# 25200056) and centrifugation for at 300 x *g* for 3 min at room temperature. After one wash with PBS (Gibco, Cat# 10010001), the cells were centrifuged as before, and the resulting cell pellet was snap frozen in liquid nitrogen and stored at –80°C. From the frozen cell pellet, total RNA was isolated using the Qiagen RNeasy Plus kit (Cat# 74134) according to the manufacturer’s protocol with the following specifications: 200 μl Buffer RLT Plus was used for cell lysis, and homogenization was carried out using QiaShredder spin columns (Qiagen, Cat# 79654) including a centrifugation at 10’000 x *g* for 3 min at 4°C. The RNA concentration and purity were determined by measuring A_260_ and A_260/280_, respectively. Subsequently, cDNA was prepared using the Transcriptor Universal cDNA Master (Roche, Cat# 05893151001) in a 20 μl reaction containing 1x Transcriptor Universal Reaction Buffer, 1x Transcriptor Universal Reverse Transcriptase and 50 ng/μl total RNA, which was incubated at 29°C for 10 min (primer annealing), at 55°C for 10 min (reverse transcription) and at 85°C for 5 min (denaturation). The newly synthesized cDNA was then diluted to 10 ng/μl and stored at –20°C. The RT-control sample consisted of RNA diluted to the same final concentration. For the qPCR assay, the Roche LightCycler® 480 System (Cat# 05015243001) in combination with 384-well plates (FrameStar, Cat# 4ti-0381/DBC, and Roche, Cat# 04729757001) was used. Each TaqMan qPCR reaction consisted of 1x LightCycler® 480 Probes Master (Roche, Cat# 04707494001), 1x Primer Probe Master (predesigned assays for IDUA, Cat# Hs.PT.58.40058589 and GAPDH, Cat# Hs.PT.39a22214836 by Integrated DNA Technologies) and 2.5 ng/μl cDNA in a final volume of 10 μl and was run in the LightCycler480 (Software version 1.5.1.62) according to the manufacturer’s guidelines. The relative mRNA levels were derived using the comparative *C*_T_ calculation method ^68^.

#### Isolation of fibroblasts from Hurler IDUA-W401X rats

Heterozygous knock-in rats were mated to produce pregnant heterozygous females. The dissected E13 embryo carcass was rinsed with cold D-PBS (Invitrogen cat# 14190-144) and dissociated in 1 ml 0.25% Trypsin-EDTA (Gibco, Cat# 25200056). After incubating in a humidified 5% CO_2_ incubator at 37°C for ∼2-3 min, 18 ml REF medium (DMEM [Gibco, Cat# 11965092] containing 10 % FCS (Bioconcept, Cat# 2-01F16-I), 100 units/ml of penicillin and 100 μg/ml of streptomycin [Invitrogen, Cat# 15140122], Gentamicin [0.5ml], Sigma Cat. #G1914 was added to the digested tissues, followed by pipetting to dissociate the tissues. Dissociated tissues were transferred to tissue culture flasks, which were cultured at 37°C in a humidified 5% CO_2_ incubator for several days until cells reached confluency. P0 rat embryonic fibroblast cells were passaged 2-3 times before compound testing.

#### Rat *in vivo* studies

CRISPR engineering and genetic analysis of the Wistar Kyoto Rat-W401X animal model was performed by GenoWay (Lyon, France). Breeding and karyotyping were outsourced to Charles River Laboratories (Wilmington, MA, USA) and Vium (San Mateo, CA, USA). Male and female animals homozygous for the IDUA W401X mutation were used at the age of 8-10 weeks (200-300 g weight) for NVS1.1 testing (IACUC: 15 DMP 067). Treatment and control groups (vehicle, WT untreated; n=5) were separately housed. Vehicle solution and compound suspensions were prepared with 0.5% (w/v) Methylcellulose, Type 1500 in aqueous solution containing 0.5% (v/v) Polysorbate 80. Formulated NVS1.1 was stored at 2-8°C and protected from light. Animal weight was measured every third day and general health was monitored daily. Pharmacokinetics of NVS1.1 was determined from tail vein and terminal blood collections. Brain tissue compound concentration was determined from collected CSF fluid. Tissues were snap frozen in liquid nitrogen and stored upon analysis.

Frozen tissue samples were pulverized with a Covaris CP02 Cryoprep device according to the manufacturer’s description. Pulverized samples were transferred in 2-or 15-ml matrix tubes, weighed and stored under ice. T-Per protein extract solution (Thermo Fisher Scientific, Cat# 78510) and protease inhibitors (Thermo Fisher Scientific, Cat# 78415) were added, vortexed and tubes were processed in a FastPrep-24^®^ Tissue and Cell homogenizer as recommended by the manufacturer. Homogenous liquid samples were stored on ice for 10 min. The process was repeated 3 times. Tissue samples were diluted to 1 g tissue/20 ml T-Per and centrifuged for 10 min. Supernatants were transferred, and all samples were stored at -80°C.

#### IDUA and GUSB enzyme assay

HEKR4 or primary fibroblast cells were seeded into black 384 well clear bottom plates (5000 cells/well) and incubated for 24 h at 37°C and 5% CO_2_ in humid atmosphere before replacing the media with NVS compound-containing media further incubation for 48 h. For 7-day treatments, the compound-containing medium was exchanged after 3 days, and cells were incubated for an additional 4 days. Cell media was removed, and cells were lysed with 3 µl cold Lysis buffer (Cell Signaling, Cat. 9803) at 4°C under shaking. The IDUA enzyme substrate (4-Methylumbelliferyl α-L-Iduronide) and GUSB enzyme substrate (4-Methylumbelliferyl -D-Glucuronide) were diluted as described by the manufacturer (Glycosynth). 5 μl of 0.4 mM of the respective substrate was added per well and incubated for 24-48 h at 37°C, 5% CO_2_. The reaction was stopped with 40 μl glycine buffer (0.5 M glycine, 0.5 M Na_2_CO_3_, pH 10.2) and substrate fluorescence (360/450nm) was measured with a PHERAstar FSX plate reader. Enzyme activity was calculated as pmol or nmol per hour per mg total protein for IDUA and GUSB, respectively. Calibration curves were derived from fluorescence substrate dilutions. Sample protein concentration was measured with a BCA kit (Thermo Fisher Scientific, Cat# 23225). For brain tissue, IDUA activity determination 20-80 μg protein samples were used, whereas for the GUSB enzymatic measurements 20 μg protein samples were sufficient. The IDUA substrate was incubated for 48 h, whereas GUSB activity was measured after 30 min substrate addition.

#### Total GAG assay

The Blyscan assay (Bicolor, Cat# B1000) is a quantitative dye-binding method for the analysis of sulfated proteoglycans and glycosaminoglycans (GAG). Total GAG from cell lysates and tissues was performed as described by the manufacturer and by others ^30^. Reference standards and reagent blanks were used for calibration curves. Lysates were mixed with 1 ml dye reagent and incubated for 30-45 min at room temperature. Precipitates were centrifuged at 10’000 x *g* for 10 min and supernatant was removed. Deposits were dissociated with 0.5 ml dissociation reagent and extensively vortexed followed by an incubation of 30 min at room temperature. Sample absorbance were measured with a microplate reader at appropriate wavelength. Plate or sample reading was done immediately. Determined GAG detection limit was 0.25 μg/sample. For brain tissue, a maximum volume of 100 μl with 300-400 μg protein/sample was used.

#### Immunoblot

HEKR4 cell pellets were resuspended in RIPA buffer (50 mM Tris pH 8, 150 mM, 5 mM EDTA pH 8.0, 1% IGEPAL CA-630, 0.5% sodium deoxycholate, 0.1 % SDS) and the lysate was isolated by centrifugation at 13’000 x *g* for 5 min at 4°C. After adjusting the protein concentrations in the lysates according to A_260_, they were supplemented with 4x NuPAGE LDS loading buffer (Thermo Fisher Scientific, Cat# NP0008) to a final concentration of 1.5x and with 25 mM DTT. Of this, samples corresponding to approx. 5×10^4^ cell equivalents were loaded per lane. Samples deriving from whole cell lysates, immunoprecipitations or polysome fractionations were resolved on 4 to 12% Bis-Tris Polyacrylamide gels (Invitrogen, Cat# NW04122BOX, Cat# WG1401BOX and Cat# WG1403BOX) in MOPS buffer. Proteins were then transferred on nitrocellulose membranes using the iBlot 2 Dry Blotting System (Invitrogen, Cat# IB21001 and IB23001). The high molecular weight protein GCN1 was transferred using the Trans-Blot Turbo Transfer System (Bio-Rad, Cat# 1704150 and Cat# 1704158). Subsequently, the membranes were blocked with 5% milk in TBS containing 0.1% tween (TBS-t) and incubated with primary antibodies (dissolved in 5% BSA in TBS-t) for 2-3 h at room temperature or overnight at 4°C. After three washes with TBS-t, the membranes were incubated with IR-Dye-conjugated secondary antibodies (dissolved in 5% milk in TBS-t) for 1 hour at room temperature. The membranes were washed again with TBS-t, dried and then scanned using the Sapphire Biomolecular Imager (Azure Biosystems). To assess eRF1 and beta-actin protein levels in compound treated fibroblasts (IDUA-W402X/Q70X Hurler patient fibroblasts or rat IDUA-W401X fibroblasts) the immunoblot was performed analogously except for the usage of horseradish peroxidase (HRP)-conjugated secondary antibodies. HRP detection was carried out using the SuperSignal™ West Femto Maximum Sensitivity Substrate. The resulting signal was measured using a BioRad ChemiDoc XRS+ imaging device, quantified and normalized to the loading control beta-actin. For each experiment, results of at least two biological replicates are shown. All primary and secondary antibodies (along with their working concentrations) that were used in this study are listed in the Key Resource Table.

#### Immunoprecipitation

For the pulldown of eRF1, one 15 cm dish of approximately 80% confluent HEKR4 PTC reporter cells was used for each condition. The cells were incubated with DMSO or 25 μM NVS1.1 for 30 min at 37°C, 5% CO_2_ in humid atmosphere, immediately washed with cold 1xPBS and kept on a bed of ice. Subsequently, the cells were scraped off the plates in 1 ml cold 1x PBS, collected by centrifugation at 500 x *g* for 5 min at 4°C, and stored at -80°C until further processing. For the immunoprecipitation, cell pellets were thawed, resuspended in IP buffer (50 mM HEPES, pH 7.3, 600 mM NaCl, 0.5% Triton X-100), and cells were then lysed by dual centrifugation using Zentrimix 380R at 1500 rpm, for 4 min at -5°C. The resulting lysate was cleared by regular centrifugation at 16’000 x *g*, for 10 min at 4°C and the protein concentration was adjusted according to A_260_. Before immunoprecipitation, 1/20 of the sample was combined with NuPAGE LDS loading buffer (Thermo Fisher Scientific, Cat# NP0008) to achieve a final concentration of 1.5x loading buffer and 25 mM DTT. The rest of the lysate was combined with 6 μg eRF1 antibody (Santa Cruz Biotechnology, Cat# sc-365686) coupled to 0.75 mg Dynabeads™ Protein G (Invitrogen, Cat# 10004D) for 1 h at 4°C under constant rotation. Subsequently, the beads were washed three times with IP buffer and the bound proteins were eluted in 1.5x LDS loading Bolt™ LDS Sample Buffer (Invitrogen, Cat# B0008) containing 25 mM DTT at 70°C for 10 min. For analysis by mass spectrometry, instead of eluting, the beads were washed three more times using IP buffer without detergent and stored at -20°C until further processing.

#### Polysome fractionation

For each condition, HEKR4 PTC reporter cells were grown on 15 cm dishes to 80% confluency and incubated with either DMSO or 25 µM NVS1.1 for 30 min. Subsequently, the cells were treated with 100 μg/ml Cycloheximide (Focus Biomolecules, Cat# 10-117) and incubated for 4 min to stop translation and stabilize ribosomes on the mRNAs. After one wash with cold 1x PBS containing 100 μg/ml Cycloheximide, the cells were harvested by scraping them off the dish in the same buffer and collecting them by centrifugation at 500 x *g* for 5 min at 4°C. The cell pellets were resuspended in 300 μl lysis buffer containing 10 mM Tris-HCl pH 7.5, 10 mM NaCl, 10 mM MgCl_2_, 1% Triton X-100, 1% sodium deoxycholate, 100 μg/ml Cycloheximide, 1 mM DTT and 0.1 u/μl RNase inhibitor (Vazyme, Cat# R301-03) and incubated on ice for 2 min with occasional vortexing to ensure complete lysis. The resulting lysates were cleared by centrifugation at 16’000 x *g* for 5 min at 4°C and transferred into a new tube. Using the BioComp gradient maker, 15-50% sucrose gradients were formed in gradient buffer (10 mM Tris-HCl pH 7.5, 100 mM NaCl, 10 mM MgCl_2_, 100 μg/ml Cycloheximide and 1 mM DTT) in ultracentrifuge tubes (Seton tubes, Cat# S7030), precooled at 4°C and balanced (+/-0.01 mg). The lysates were loaded onto the sucrose gradients and centrifuged at 40’000 rpm for 2 h at 4°C using the SW 41 Ti rotor (Beckman Coulter, Cat# 331362). Subsequently, the gradients were fractionated into 1.5 ml tubes using the BioComp piston fractionator (volume displaced 0.143 ml/mm, scan speed 0.30 mm/sec, distance 3.92 mm/fraction, 0.559 ml/fraction) and the A_260_ profile was recorded. The retrieved fractions were then stored at -80°C. For protein analysis, the fractions were mixed with 1 volume of acetone and 0.1 volumes of 100% TCA, incubated overnight at -80°C, and the precipitated proteins were collected by centrifugation at 16’000 x *g* for 5 min at 4°C. Pellets were washed three times with ice-cold acetone (centrifugation as before), dried for 20 min in a SpeedVac and stored at -80°C. For immunoblot analysis, the protein pellet of every odd-numbered fraction was dissolved in NuPAGE LDS loading buffer (Thermo Fisher Scientific, Cat# NP0008) to a final concentration of 1.5x and 25 mM DTT. 7% of the collected fraction was loaded per lane. For analysis by label-free mass spectrometry, 50% of two heavy polysome fractions (between fractions 16 and18, depending on the individual replicate) were pooled and precipitated as described above.

#### SILAC experiment in HEK293T cells to assess whole proteome changes under NVS1.1 treatment

HEK293T cells were metabolically labeled with heavy or light amino acids and treated for 24 h with 1 µM NVS1.1 or DMSO with swapping labels. Cells were harvested and pellets were resuspended in a buffer of 50 mM HEPES, 150 mM NaCl, 1.5 mM MgCl_2_, and 1% SDS and lysed to completion by sonication with a probe tip sonicator. Samples were mixed in a 1:1 ratio according to protein concentration determined with the 660 nm Protein Assay (Pierce). A total protein input of approximately 20 µg was used per replicate experiment. Lysates were reduced with 10 mM DTT (Indofine Chemicals) at 30°C for 30 min, followed by alkylation with 25 mM iodoacetamide (Sigma-Aldrich) for 30 min at room temperature in the dark. Following SDS removal using detergent removal spin columns (Pierce, Cat# 87777), samples were digested overnight at 37ºC with trypsin (Pierce, Cat# 90057) at a trypsin/protein ratio of 1:60, acidified with 5% formic acid and lyophilized. After resolubilization in separation buffer A (4% 5 mM ammonium formate, pH 10), in tryptic peptides were separated by high pH reversed-phase HPLC using a Dionex 3000 fitted with a 6.4 × 150 mm Zorbax C18 extend column with a flow rate of 1 mL/min. Eighty 1 ml fractions were collected throughout the segmented gradient (0-8% B/7 min, 8-27% B/38 min, 27-31% B/4 min, 31-39% B/8 min, 39-60% B/7 min, 60-0% B/20 min, separation buffer B: 96% acetonitrile, 4% 5 mM ammonium format, pH 10). After pooling to 17 fractions based on the UV absorption profile to achieve comparable peptide content per sample, the resulting fractions were analyzed by nanoLC-MS/MS. A nano LC column was prepared by creating a pulled tip with a P-2000 laser puller (Sutter Instruments) and packing the 75 µm ID fused silica with C18 material (Dr Maisch Reprosil Pur120, C18AQ 3 µm) to a length of 15 cm and fitted to the Eksigent nano \LC (AB SCIEX). The eluted peptides were reconstituted in 100 μl 2% acetonitrile in 0.1% formic acid (LC-MS Buffer A) and 25 μl was injected to the mass spectrometer (hybrid LTQ-Orbitrap-Velos-Elite, Thermo Fisher Scientific) using a trapping column (1 cm Michrom Magic C18AQ, 5 μm), washed for 20 min and then switched in-line with the analytical column. The peptides were eluted with a gradient of 3% LC-MS buffer B (70% acetonitrile in 0.1% formic acid) to 45% B in 80 min (0.5% B/min) delivered at a flow rate of 300 nl/min and using a top 20 CID analysis method with a dynamic exclusion set to 2. Data were analyzed with MaxQuant version 1.3.0.5 (Andromeda search engine with MaxQuant quantitation), searched against the UniProt Human database (V9 plus typical lab contaminants) with the addition of MaxQuant-generated reversed database to calculate false discovery rates. Database search criteria required full tryptic cleavage and allowed for up to 2 missed cleavages with oxidized methionine, and N-terminal acetylation as variable modifications and carbamidomethylation of cysteine as static modification. The results were visualized using GraphPad Prism.

#### Determination of amino acid inserted at stop codon

##### SDS-PAGE separation and in-Gel digestion

HEKR4 cells stably expressing IDUA-W402X with different stop codons (UAA, UAG or UGA) treated with 2 μM NVS1.1, 5 NVS2.1 or 14.4 mM Paromomycin for 24 hours. The cells were then harvested, washed twice in PBS and after centrifugation at 250 x g for 5 min at 4°C resuspended in cell lysing buffer (Cell Signaling Cat# 9803, containing protease inhibitor, Thermo Fisher Scientific, Cat# 78415). Cell lysates were incubated on ice for 45 min with occasional mixing. Cell lysates were supplemented with LDS loading buffer containing 100 mM DTT. In parallel, a sample containing recombinant Human iduronidase (Gentaur, Cat# MBS717919) was prepared as reference. Separation of proteins was performed by SDS-PAGE using a NuPage Bis-Tris 4-12% gradient minigel (Invitrogen, Cat# NP0335) as described by the manufacturer. Approximately 0.3-0-8 μg of protein was applied to each gel lane. Three bands (approximately 1 mm) were typically excised between 70-80 kDa and further subjected to in-gel digestion In-gel digestion was performed as previously described ^69^ using a perforated 96-well microtiter plate format (CB080; Proxeon; DEN). Endoproteinase AspN (Roche Diagnostic, Cat# 11054589001) digestion of the gel pieces was performed in a 50 mM ammonium bicarbonate buffer (typically 240 ng of enzyme per sample)

##### NanoLC-MS/MS - Parallel Reaction Monitoring (PRM)

Separation of peptides produced by AspN digestion was performed on a ICS3500 nano-UHPLC system (Thermo Fisher Scientific/Dionex, Germering, DE)) employing a 75 μm x 150 mm Easy spray column packed with C-18 reverse phase (Thermo Fisher Scientific, Cat# ES901) and a trapping column (Thermo Fisher Scientific, Cat# 164197). The column and trapping columns were kept at 40°C. Sample volumes of 5 μl were injected onto the trapping column. UPLC was controlled by Chromeleon software (Thermo Fisher Scientific). Eluent A was water containing 0.1% TFA. Eluent B was a 1:9 mixture of water: acetonitrile containing 0.09% TFA. A gradient from 20% B to 90% B was run in 60 min. The flow rate was typically 300 nl/min. The mass spectrometer was a QExactive (Thermo Fisher Scientific) equipped with Easy-spray ESI source. Parallel reaction monitoring (PRM) was performed on the doubly charged ions from the target peptides of interest. ^70^. A list of 20 different target Asp-N digestion peptides containing the amino acid in position 402 of IDUA was monitored. Retention time and most abundant fragment information were obtained from LC-MSMS experiments using synthetic peptides from sequence D_397_-L_412_ (20 peptides each containing a different amino acid in position 402; see list in accessory records) Data were acquired and processed using the Excalibur software (Thermo Fisher Scientific). Semi-quantitative data were obtained from extracted ion intensities of peptide fragments

#### Label-free mass spectrometry for whole proteome and ubiquitination analysis

After immunoprecipitations, proteins associated with the Dynabeads were resuspended in 8 M urea, 50 mM Tris-HCl pH 8, reduced at 37°C for 30 min with 0.1 M DTT, 50 mM Tris-HCl pH 8, alkylated at 37°C for 30 min in the dark with 0.5 M iodoacetamide (IAA), 50 mM Tris-HCl pH 8, diluted with 4 volumes of 20 mM Tris-HCl pH 8 2 mM CaCl_2_ prior to overnight digestion with 100 ng sequencing grade trypsin (Promega, Cat# V5111) at room temperature. Samples were then centrifuged, the magnetic beads trapped by a magnet, and the peptides-containing supernatant was collected.

The peptides were analyzed by liquid chromatography (LC)-MS/MS (PROXEON coupled to a QExactive HF mass spectrometer, Thermo Fisher Scientific) with three injections of 5 μl digests. Peptides were trapped on a μPrecolumn C18 PepMap100 (5μm, 100 Å, 300 μm × 5 mm, Thermo Fisher Scientific, Reinach, Switzerland, Cat# 160454) and separated by backflush on a C18 column (5 μm, 100 Å, 75 μm × 15 cm, C18, NYKKYO, Cat# NTCC-360/75-3-155) by applying a 60-min gradient of 5% acetonitrile to 40% in water, 0.1% formic acid, at a flow rate of 350 nl/min. The Full Scan method was set with resolution at 60,000 with an automatic gain control (AGC) target of 1E06 and maximum ion injection time of 50 ms. The data-dependent method for precursor ion fragmentation was applied with the following settings: resolution 15,000, AGC of 1E05, maximum ion time of 110 ms, mass window 1.6 m/z, collision energy 28, under fill ratio 1%, charge exclusion of unassigned and 1+ ions, and peptide match preferred, respectively.

For whole proteome analyses after polysome fractionations, cell pellets were re-suspended in 12 μl lysis buffer (8M Urea, 100 mM Tris-HCl pH 8, and a 1:10 dilution of 2 μl of the samples was used to measure protein concentration by Qubit Protein Assay (Invitrogen, Cat# Q33211). The remaining 10 μl were reduced, alkylated, and digested with LysC (Promega, Cat# VA117A) for 2 h at 37°C, followed by 100 ng Trypsin (Promega, Cat# V5111) overnight digestion at room temperature. The digests were analyzed by liquid chromatography on a Dionex, Ultimate 3000 (Thermo Fisher Scientific) coupled to a LUMOS mass spectrometer (Thermo Fisher Scientific) with two injections of 500 ng peptides.

The samples were loaded in random order onto a pre-column (C18 PepMap 100, 5 μm, 100 A, 300 μm i.d. x 5 mm length) at a flow rate of 10 μl/min with solvent C (0.05% TFA in water/acetonitrile 98:2). After loading, peptides were eluted in back flush mode onto a homemade pack C18 CSH Waters column (1.7 μm, 130 Å, 75 μm × 20 cm) by applying a 90-min gradient of 5% to 40% acetonitrile in water, 0.1% formic acid, at a flow rate of 250 nl/min.

Data acquisition used the data dependent mode, with precursor ion scans recorded in the orbitrap with resolution of 120’000 (at m/z=250) parallel to top speed fragment spectra of the most intense precursor ions in the linear trap for a cycle time of max. 3 seconds.

Peptides and proteins were searched and quantified using FragPipe version 1.8 ^71-73^. A closed search was performed with the Swissprot ^74^ human database containing isoforms (release June 2022), to which reverse decoys and common contaminants were added. Fragment mass tolerance was set to 20 ppm for IP samples and to 0.4 Da for samples from lysed cells, respectively. Search enzyme was set to “strict trypsin” and allowed missed cleavages to 3. Carbamidomethylation was set as a fixed modification on cysteine, and the following variable modifications were enabled: methionine oxidation, ubiquitination residue on Lysine and protein N-terminal acetylation. Philosoper’s peptide and protein prophet ^71^ were selected, respectively, for filtering and protein inference with default parameters, and IonQuant ^73^ for quantification. Match between runs was enabled, with top runs set to 2 for samples from beads and 3 for samples from cell pellets.

Differential protein expression was determined as follows: Protein groups with only 1 peptide evidence and the common contaminants were removed. Imputation was performed if there were at least 2 detections in at least one group of replicates. If there was at most 1 non-zero value in the group for a protein, then the remaining missing values were imputed by drawing values from a Gaussian distribution of width 0.3 times the sample standard deviation and centered at the sample distribution mean minus 2.5 times the sample standard deviation. Any remaining missing values were imputed by the Maximum Likelihood Estimation ^75^ method. Differential expression by moderated t statistics between the two groups and significance evaluation was performed as previously described ^76^. The relative modification degree for ubiquitination residues was obtained by first summing the contributing intensities to each ubiquitination site in each sample, then dividing this quantity by the protein intensity, and averaging the result across the replicates.

