## Supplemental Figures and Table for "Drug-induced eRF1 degradation promotes readthrough and reveals a new branch of ribosome quality control"

### Figure S1

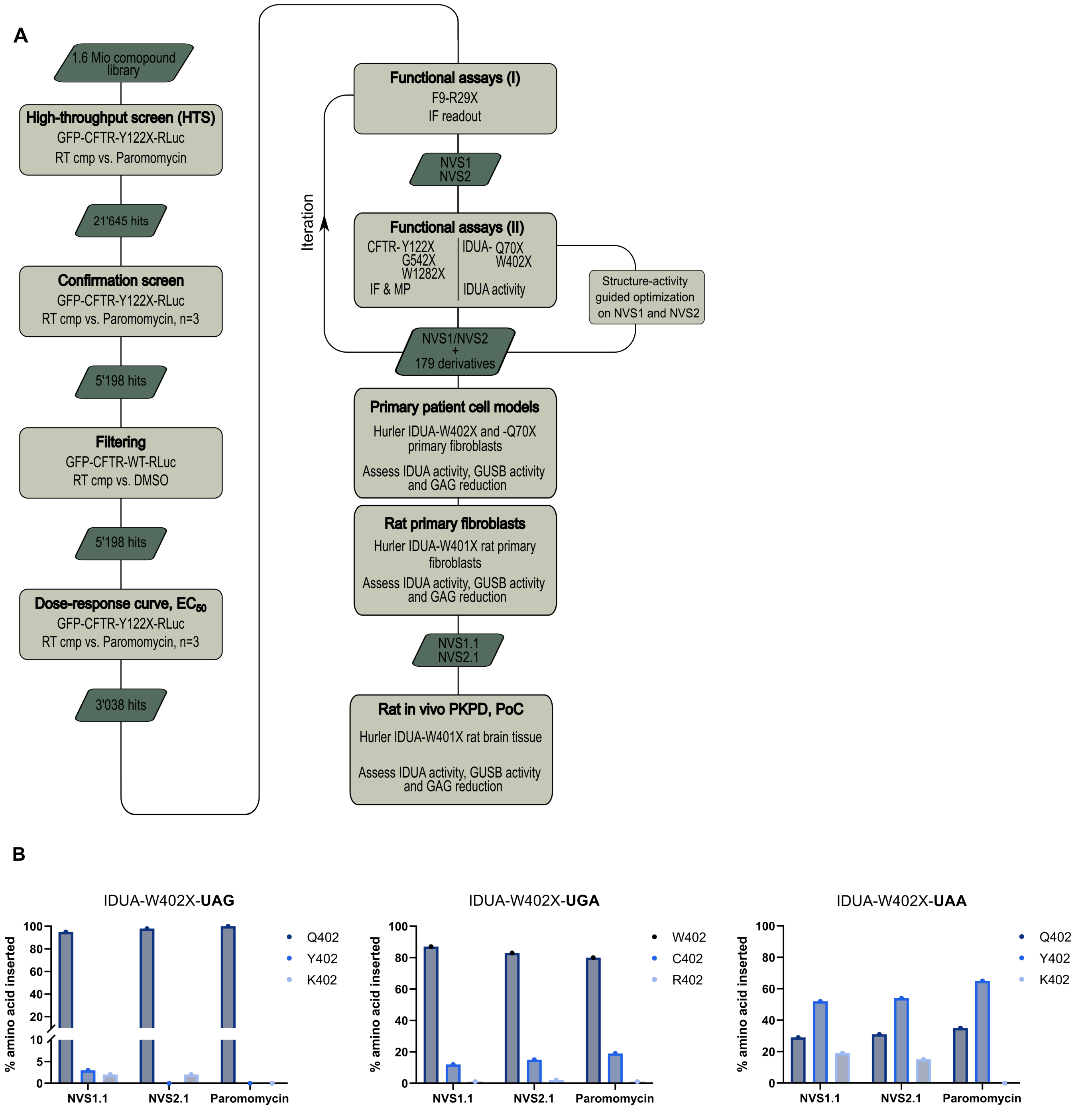


**Figure S1, related to Figure 1.**
(A) Flowchart depicting the lead compound identification and triaging of positive hits. Abbreviations: Half maximal effective concentration ($EC_{50}$), immunofluorescence (IF), membrane potential (MP), proof of concept (PoC).
(B) Quantitative Nano-LC-MS/MS to determine the incorporated amino acid at each of the three TCs in recombinant W402X-IDUA expressing HEKR4 cells in response to NV1.1, NVS2.1 or Paromomycin.

### Figure S2

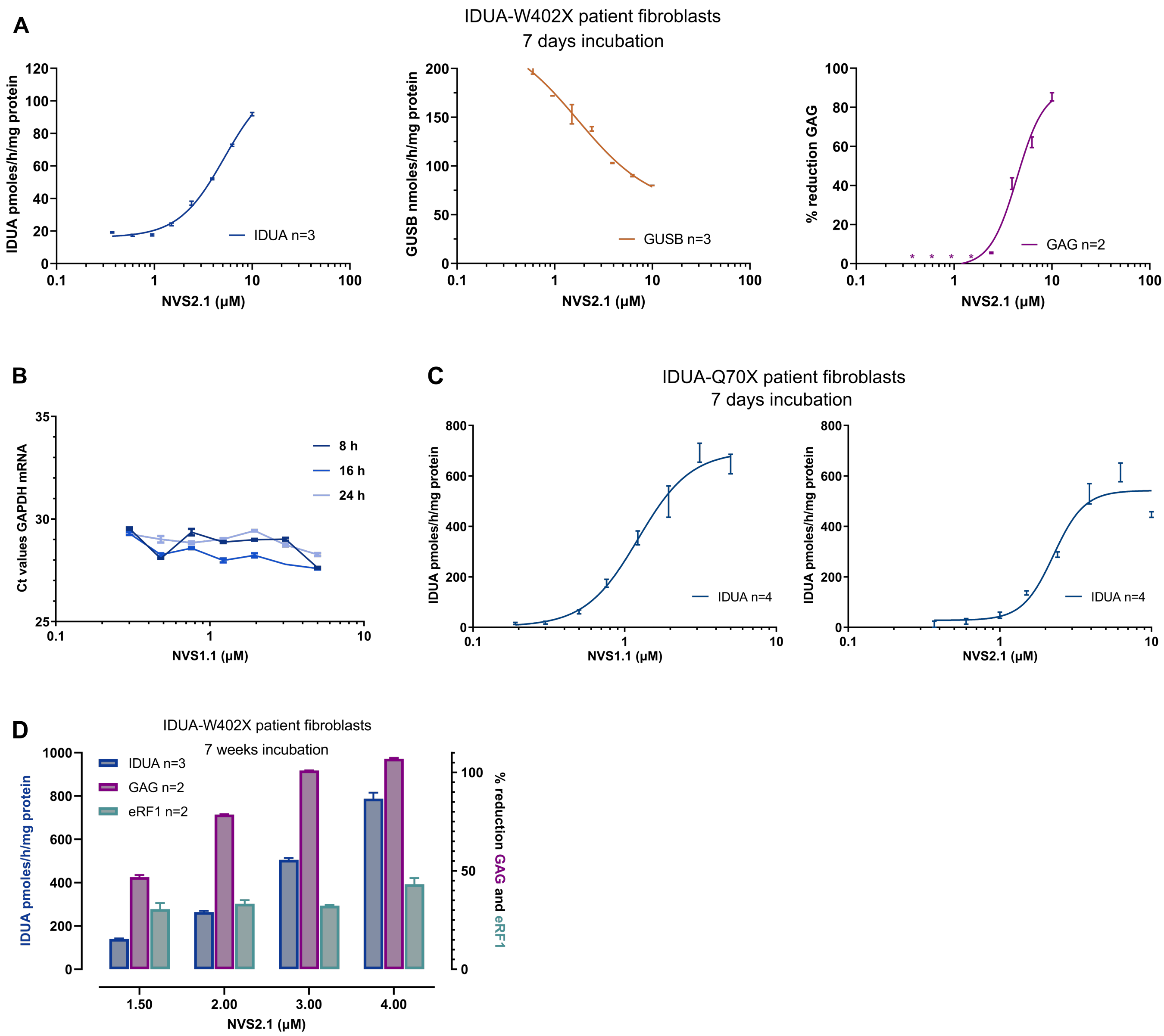


**Figure S2, related to Figure 2.**

(A) Primary fibroblasts deriving from patients homozygous for the IDUA W402X mutation were treated for 7 days with the indicated concentrations of NVS2.1. Subsequently, α-L-iduronidase (IDUA) and β-glucuronidase (GUSB) activities were assessed by measuring the conversions rates of the surrogate substrates 4-Methylumbelliferyl-α-L-iduronide (pmol/h/mg protein) and 4-Methylumbelliferyl-β-D-glucuronide (nmol/h/mg protein), respectively. Total glycosaminoglycan (GAG) levels were determined by a colorimetric assay and normalized to the vehicle-treated control sample. Asterisks denote GAG measurements below the assay detection limit (resulting in negative values) that were removed from the dataset.

(B) RT-qPCR assay showing the GAPDH mRNA Ct (cycle threshold) values upon treatment of the Hurler primary fibroblasts with different concentrations of NVS1.1 for 8 – 24 hours.

(C) Primary Hurler patient fibroblasts homozygous for the IDUA Q70X mutation were cultured for 7 weeks in the presence of NVS1.1 or NVS2.1 (compound exchange every 3rd day) and α-L-iduronidase enzymatic activity was assessed as in (A)

(D) Primary Hurler IDUA-W402X fibroblasts were cultured in different concentrations of NVS2.1 with medium and compound exchange every 3rd day. After 7 weeks, the IDUA activity (left y-axis) and total GAG levels (right y-axis) were determined as in (A). The eRF1 protein abundance (right y-axis) was assessed by immunoblot (using beta-actin as a loading control) and is expressed relative to the DMSO-treated control condition.

#### Figure S3

**A**

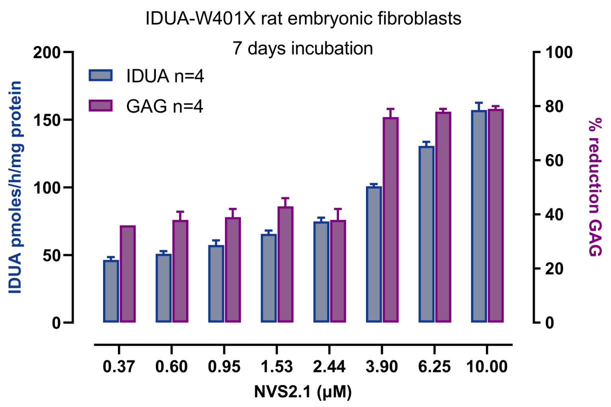


**B**

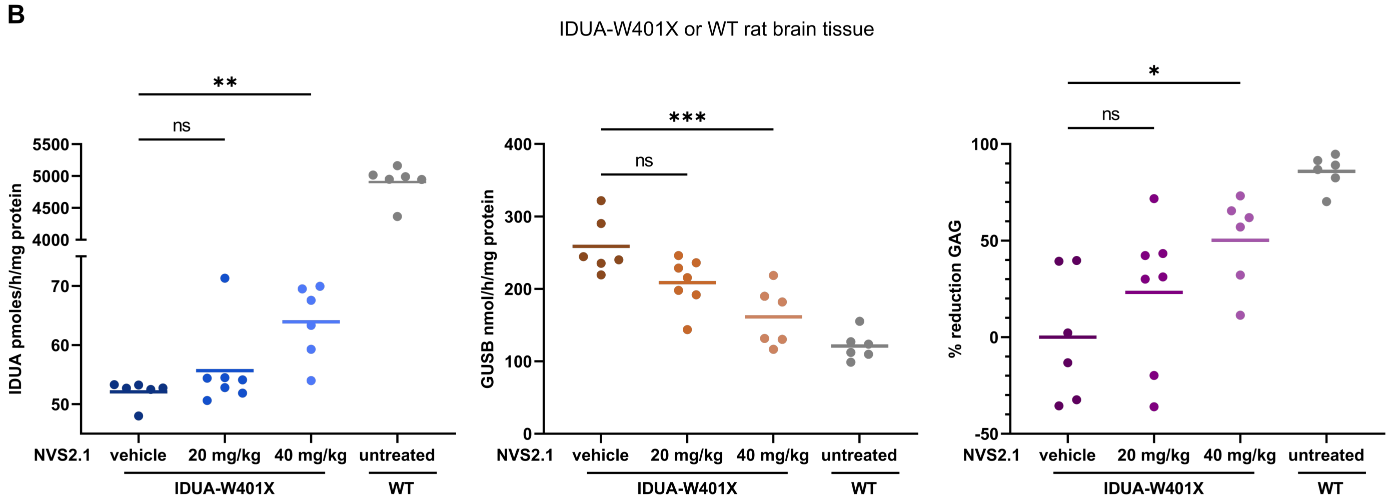


**Figure S3, related to Figure 3.**

(A) Freshly isolated rat fibroblasts of a Hurler animal model homozygous for IDUA-W401X were treated for 7 days with different concentrations of NVS2.1. The IDUA enzyme activity and total GAG levels were determined as in Figure S2A.

(B) Rats homozygous for the IDUA W401X mutation were treated with 20 mg/kg (n=7) or 40 mg/kg (n=6) NVS2.1, or with the vehicle formulation only (n=7) for 14 days. A similar number of untreated animals with WT IDUA was analyzed for comparison. After scarifying the animals, the brain tissue was extracted and the activities of IDUA and GUSB along with the total GAG levels were determined as in Figure 2A. In each graph (IDUA, GUSB, GAG), one dot represents the average of four technical replicates carried out with brain tissue from one animal. *P* values were calculated by one-way ANOVA followed by Dunnet's multiple comparisons test. *P* values of ≤ 0.05, p≤ 0.01, p≤ 0.001 and >0.05 are depicted as *, **, ***, and ns (non-significant), respectively.

### Figure S4

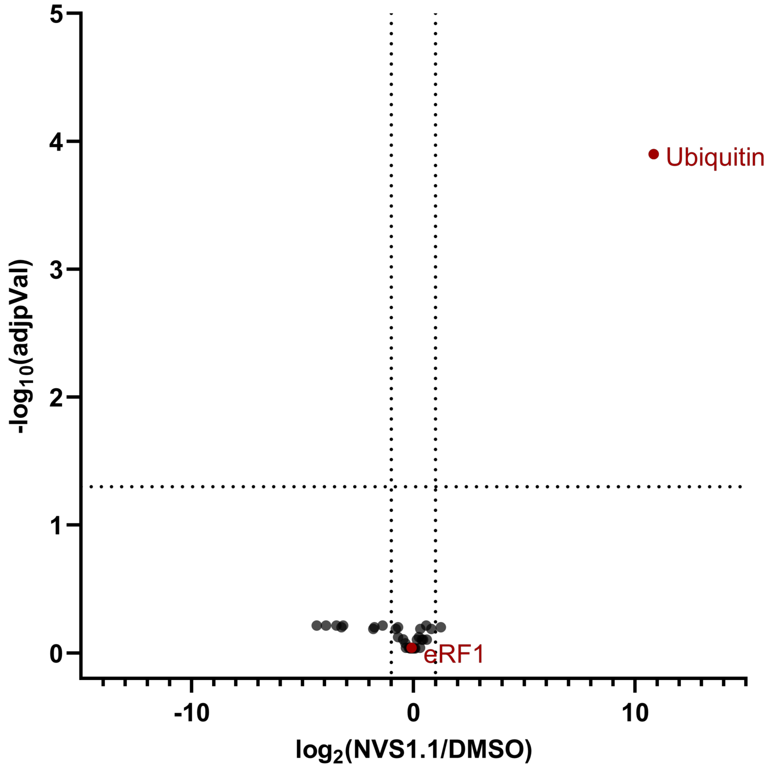


**Figure S4, related to Figure 4.**
Label-free mass spectrometry analysis depicting the fold change of identified proteins upon treatment with NVS1.1 ($log_2$(NVS1.1/DMSO), x-axis), versus its statistical significance ($log_{10}$adjp-value, y-axis) of the immunoprecipitates of Figure 4A.

### Figure S5

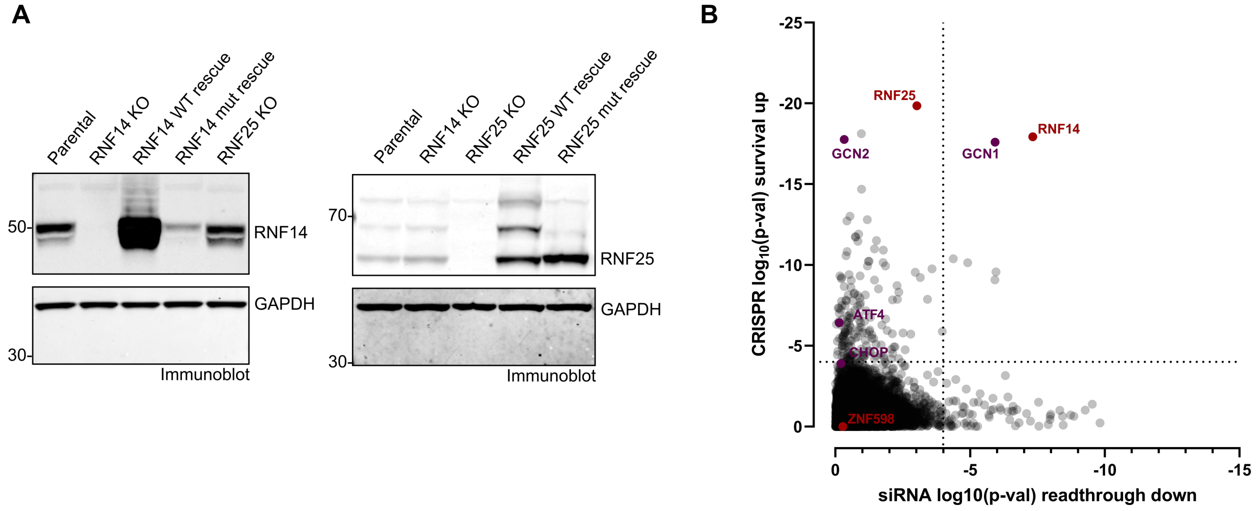


**Figure S5, related to Figure 5**

(A) Detection of the RNF14 and RNF25 protein levels in the knockout and rescue cell lines by immunoblotting. GAPH served as a loading control.

(B) Combined results of a genome-wide siRNA screen scoring for reduced NVS2.1-mediated readthrough and a CRISPR knockout screen enriching for genes leading to NVS2.1 resistance. X-axis: $Log_{10}$ p-values depicting the significance of the reversion of NVS2.1-induced readthrough in CFTR-Y122X-Rluc reporter gene expressing HEKR4 PTC cells for the knockdown of each of the 19’300 tested genes. For each gene knockdown (8 siRNAs per gene), Rluc activity was normalized to the luminescence signal of a non-targeting siRNA ($log_2$FC) and the differential activity between the two treatment conditions (NVS2.1 $IC_{80}$ vs. DMSO) was determined. The gene significance was calculated for the differential activity of each gene knockdown using the RSA statistical test (redundance siRNA activity; Konig et al., *Nat Methods* 4(10):847-9, 2007). Y-axis: $Log_{10}$ p-values depicting the significance of the reversion of NVS2.1-induced toxicity in a cell survival assay for the knockout of each of the 19’300 tested genes. Differential representation of each sgRNA in NVS2.1 $IC_{80}$ and untreated library-infected cell populations was determined as surrogate of difference in cell proliferation. The gene significance was calculated for the differential representation of each sgRNA set (5 sgRNAs per gene) using the RSA statistical test. For both screens, the significance thresholds were determined by randomizing the gene labels before running the RSA tests. A $log_{10}$(p-Val) <-4 threshold (dotted lines) limited false positives to ~5%.

### Figure S6

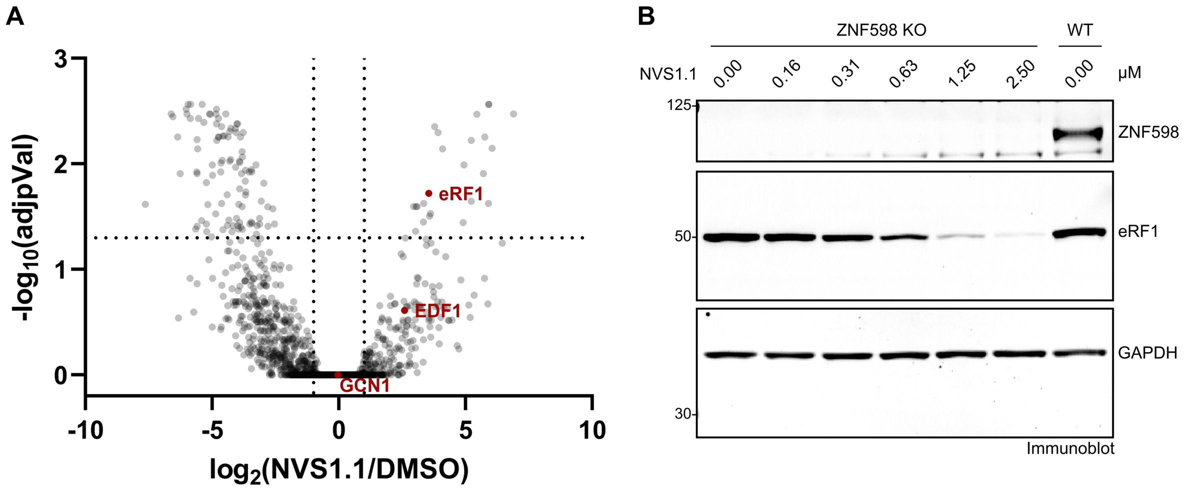


**Figure S6, related to Figure 6.**
(A) The relative abundances of detected proteins in the mass spectrometry dataset from Figure 6C were determined and are depicted as volcano plot showing the fold change $\log_2$(NVS1.1/DMSO) on the x-axis and the adjusted p-value ($-\log_{10}$(adjpVal)) on the y-axis.
(B) HEK Flip-In T-REx cells containing a ZNF598 knockout (ZNF598 KO) were incubated with NVS1.1 for 6 hours and analyzed for eRF1 by immunoblot using GAPDH as normalizer. In addition, lysate of untreated HEK cells was loaded to confirm the depletion of ZNF598 on the immunoblot.

**Table S1**

| Oligonucleotides | | |
|---|---|---|
| Name | Sequence 5'-3' or identifier | Supplier |
| IDUA TaqMan assay | Cat# Hs.PT.58.40058589 | Integrated DNA technologies |
| GAPDH TaqMan assay | Cat# Hs.PT.39a22214836 | Integrated DNA technologies |
| sgRNA_RNF25_fwd | accgACCCTCTAGATGTAGTGAAA | - |
| sgRNA_RNF25_rev | aaacTTTCACTACATCTAGAGGGT | - |
| sgRNA_RNF14_fwd | accgGTGCAGGTTGACCTACCATG | - |
| sgRNA_RNF14_rev | aaacCATGGTAGGTCAACCTGCAC | - |
| 5644 | AATGATACGGCGACCACCGAGATCTACACTCGATTTCTTGGCTTT ATATATCTTGTGGAAAGGA | Integrated DNA technologies |
| INDEX | CAAGCAGAAGACGGCATACGAGATXXXXXXXXXXGTGACTGGAG TTCAGACGTGTGCTCTTCCGATC | Integrated DNA technologies |
| P5 | AATGATACGGCGACCACCGAGA | Integrated DNA technologies |
| P7 | CAAGCAGAAGACGGCATACGA | Integrated DNA technologies |
| 5645 | TCGATTTCTTGGCTTTATATATCTTGTGGAAAGGACGAAACACCG | Integrated DNA technologies |
